# Design of multimodal antibiotics using deep learning

**DOI:** 10.1101/2024.12.20.629780

**Authors:** Angela Cesaro, Fangping Wan, Marcelo D. T. Torres, Cesar de la Fuente-Nunez

## Abstract

The rise of antimicrobial resistance has rendered many treatments ineffective, posing serious public health challenges. Intracellular infections are particularly difficult to treat since conventional antibiotics fail to neutralize pathogens hidden within human cells. However, designing molecules that penetrate human cells with preserved antimicrobial activity has historically been a major challenge. Here, we introduce ApexDuo, a multimodal artificial intelligence (AI) model for generating peptides with both cell-penetrating and antimicrobial properties. From a library of 50 million AI-generated compounds, we selected and characterized several candidates. Our lead, Turingcin-46, penetrated mammalian cells and reduced intracellular *Staphylococcus aureus* and *Listeria monocytogenes*. In mouse models of skin abscess and peritonitis, Turingcin-46 reduced bacterial loads by up to two orders of magnitude. In sum, ApexDuo generated multimodal peptide antibiotics, opening new avenues for designing molecules with more than one function.

## Introduction

Antimicrobial resistance is a critical global health threat, with nearly three million infections caused by resistant pathogens occurring annually in the United States alone, resulting in over 35,000 deaths and treatment costs exceeding $4.6 billion ^1,2^. As pathogenic bacteria evolve diverse resistance mechanisms, they can also employ strategies to evade both host immune defenses and conventional antibiotics. Among these strategies is intracellular pathogenicity, wherein certain bacterial species, such as *Staphylococcus aureus*, invade mammalian cells, replicate intracellularly, and subsequently re-emerge to infect new tissues. This process can lead to severe conditions, including sepsis ^3–5^. Addressing this challenge requires novel therapeutics that combine multiple and complementary biological activities.

Traditional drug discovery strategies have proven time-consuming, costly, and increasingly inadequate for addressing the escalating resistance problem. Moreover, these methods predominantly target extracellular bacteria, failing to address the pressing need for therapies capable of eradicating intracellular pathogens ^6,7^.

Here, we introduce ApexDuo, a deep learning model designed to generate peptide candidates with both cell-penetrating and antimicrobial properties. Cell penetrating peptides (CPPs) facilitate the translocation of biomolecules, including therapeutics, into mammalian cells, while antimicrobial peptides (AMPs) offer broad-spectrum antimicrobial activity against diverse pathogens and may modulate immune responses ^8–10^. Our objective was to create novel, multimodal molecules capable of penetrating host cells and reducing intracellular bacteria, thereby addressing critical gaps in the treatment of persistent intracellular infections.

Using ApexDuo, we generated a library of 50 million peptide candidates, synthesized and experimentally tested 60 sequences —collectively termed Turingcins in honor of Alan Turing. Among these, one peptide, Turingcin-46, exhibited remarkable efficacy in suppressing intracellular *S. aureus* infections *in vitro*. Finally, we provide evidence that Turingcin-46 displays anti-infective activity in two different mouse models.

## Results

### Overview of ApexDuo

To design peptides with dual cell-penetrating and antimicrobial properties, we leveraged a deep generative model, specifically a maximum mean discrepancy variational autoencoder (MMD-VAE), to produce novel peptide sequences. We then applied deep learning-based predictors to assess both cell-penetrating and antimicrobial activities, identifying candidates with the desired multimodal profiles **(Figure 1A)**. From ∼50 million peptides generated, we applied stringent filtering criteria to select 60 top-ranked compounds for synthesis and characterization **(Figure S1A**, see also “**Peptide selection**” subsection of the **Methods)**. These criteria included excluding peptides with high median MIC values, low cell-penetration probabilities, duplicates, and peptides already present in in-house datasets and public databases, as well as minimizing sequence redundancy. After applying these filters, 52 soluble peptides—collectively termed Turingcins—were selected for experimental testing **(Table S1)**.

**Figure 1.**
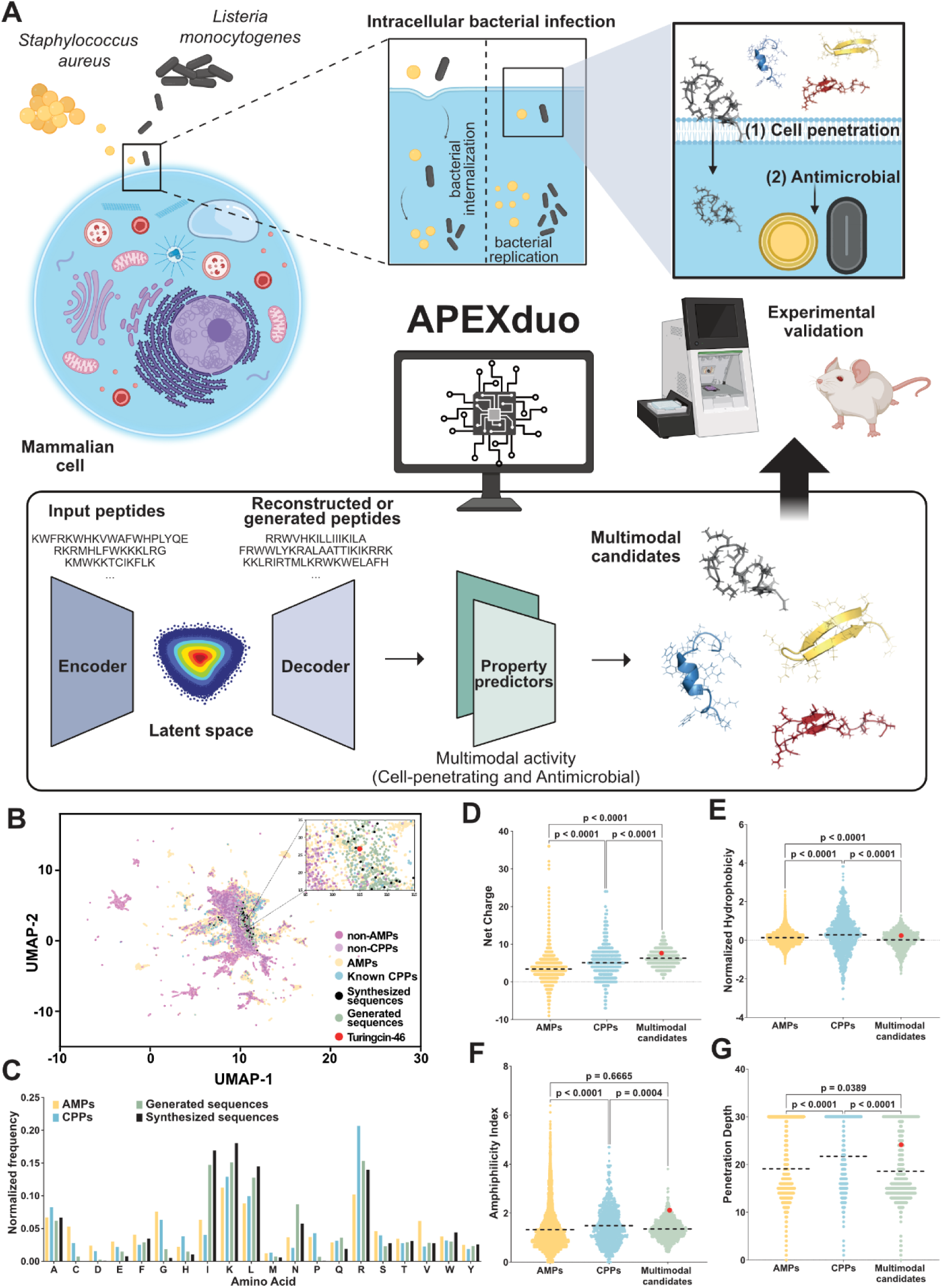
Multimodal peptide generation using deep learning. **(A)** A deep learning workflow capable of generating peptide candidates that can penetrate eukaryotic cells and eliminate intracellular *S. aureus* and *L. monocytogenes.* A variational autoencoder (VAE) was used to generate novel peptide sequences. The resulting candidates were then filtered using predictive deep learning models to select those with high antimicrobial and cell- penetrating potentials for further experimental validation. The peptide structures shown were rendered using the PyMOL Molecular Graphics System (version 3.0, Schrödinger, LLC). Figure created with BioRender.com. **(B)** Distribution of different peptide classes in ESM2-derived sequence space. Sequences from publicly available AMPs and CPPs, as well as multimodal candidates generated by ApexDuo, were embedded into a 2D space after dimensionality reduction via a sequence similarity matrix. This analysis reveals that the newly generated multimodal candidates occupy a region distinct from non-AMP/CPP sequences, yet overlap with the known AMP/CPP space, suggesting the potential for dual functionality. **(C)** Comparison of amino acid frequencies in Turingcins (both generated and synthesized) against publicly available AMPs and CPPs. The multimodal peptides show a unique compositional profile, including a relative enrichment of lysine, isoleucine, leucine, and asparagine residues. **(D–G)** Comparative analysis of physicochemical properties among the multimodal candidates, AMPs, and CPPs. Properties include **(D)** net charge, **(E)** normalized hydrophobicity, **(F)** amphiphilicity index, and **(G)** penetration depth. Normalized hydrophobicity influences peptide-membrane interactions, net charge modulates electrostatic attractions with negatively charged bacterial membranes, amphiphilicity index correlates with the peptide’s mechanism of membrane engagement, and penetration depth reflects how deeply the peptide integrates into lipid bilayers. All physicochemical properties of the peptides were obtained using the DBAASP server ^39^, and the hydrophobicity scale used was the Eisenberg and Weiss scale. Turingcin-46, the most promising multimodal candidate identified in this study, is indicated with a red For panels **D–G**, statistical significance was determined via two-tailed t-tests followed by Mann–Whitney tests; p values are shown on the graphs. The dashed line within each distribution denotes the mean for that group.

### Composition and physicochemical features of multimodal candidates

To investigate the distribution and similarity of the generated multimodal candidates, we used first token embeddings from the pretrained protein large language model ESM2 to represent peptides from various sources, including non-AMPs/CPPs, AMPs/CPPs, generated peptides, and synthesized peptides. These representations were plotted in 2D using Uniform Manifold Approximation and Projection (UMAP), revealing that the generated multimodal candidates overlapped with known AMPs/CPPs. This overlap suggests their potential to exhibit both antimicrobial and cell-penetrating properties **(Figure 1B)**.

An analysis of amino acid composition revealed distinctive features of Turingcins. These peptides showed slight enrichments in isoleucine, lysine, leucine, and asparagine frequencies, contributing to their higher net positive charge relative to common AMPs and CPPs, which generally exhibit lower lysine levels. Conversely, Turingcins displayed reduced levels of cysteine, aspartic acid, glutamic acid, and glycine, as well as lower arginine content compared to CPPs, and lower valine content compared to AMPs. Notably, the synthesized and validated Turingcins lacked cysteine, aspartic acid, and proline residues altogether **(Figure 1C)**.

Regarding physicochemical properties, the multimodal candidates demonstrated lower normalized hydrophobicity values compared to known AMPs and CPPs but exhibited intermediate amphiphilicity index values—an important trait for peptides designed to penetrate biological membranes **(Figures 1D-G)**. This suggests their membrane interactions are less intense than those of highly amphiphilic peptides. The Turingcins showed intermediate values for amphiphilicity, hydrophobicity, and disordered conformation propensity, bridging the characteristics of AMPs and CPPs. However, their net charge, hydrophobic residue angle, and isoelectric point aligned more closely with CPPs, while other features more closely resembled AMPs **(Figures 1D-G and S1B-I).**

These distinct physicochemical and compositional features set Turingcins apart from previously described peptide classes, highlighting their unique potential as multimodal therapeutics.

### *In vitro* antimicrobial activity of multimodal candidates

The antimicrobial efficacy of the designed Turingcins was initially evaluated against twelve clinically relevant bacterial strains, encompassing both Gram-positive and Gram-negative species, with a focus on members of the ESKAPEE group—recognized by the World Health Organization as major public health threats ^11^ **(Figure 2B)**. A remarkable 90.38% (47/52) of Turingcins exhibited activity against at least one pathogen with MIC values (minimal inhibitory concentrations resulting in complete inhibition of microbial growth) ranging from 0.78 to 50 μmol L^-1^. Among these, 15 peptides demonstrated activity against *S. aureus* ATCC 12600.

**Figure 2.**
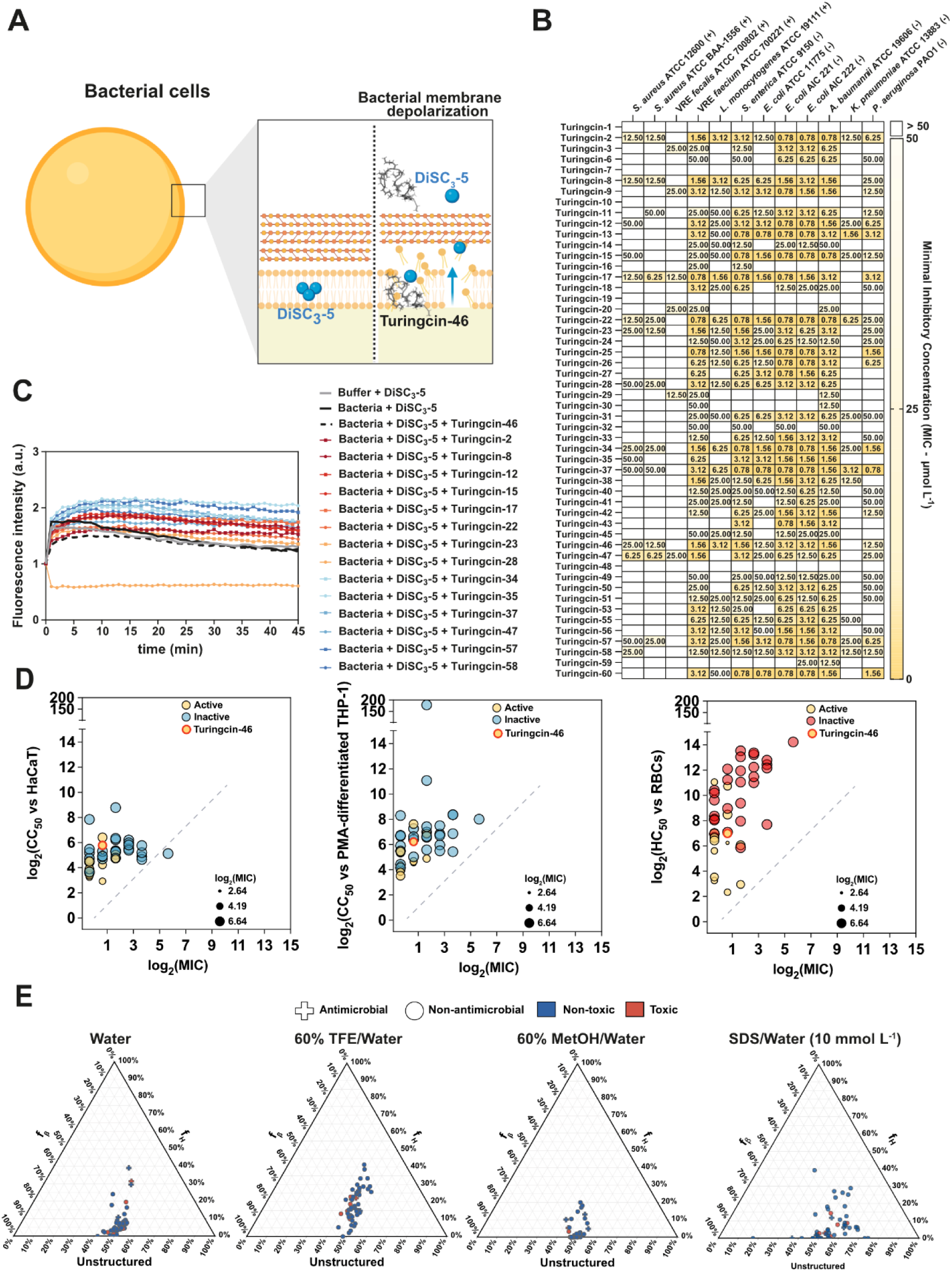
Antimicrobial activity, cytotoxicity, and secondary structure profiles of Turingcins. **(A)** Schematic overview of how Turingcins exert antimicrobial activity against *S. aureus*, including the induction of membrane depolarization. **(B)** Heat map showing the minimal inhibitory concentration (MIC) values of active Turingcins generated by ApexDuo against clinically significant Gram-positive (+) and Gram-negative (-) bacterial strains, including antibiotic-resistant isolates. Cells were treated with serial dilutions of each peptide (0.78–50 µmol L^-1^) at 37 °C, and bacterial growth was measured after 24 hours via optical density at 600 nm. MIC values indicate the lowest concentration that completely inhibited growth, reported as the mode of two independent replicates. **(C)** Relative fluorescence changes of *S. aureus* membranes treated with Turingcins in the presence of the voltage-sensitive dye DiSC3-5, indicating peptide-induced membrane depolarization. All experiments were performed in three independent replicates (see also **Figure S2A**) **(D)** Scatterplots correlating antimicrobial activity (lowest MIC observed across all tested pathogens) with cytotoxicity [CC50 for human keratinocytes (HaCaT), PMA-differentiated human monocytes (THP-1), and HC50 for human red blood cells (RBCs)]. CC50 and HC50 values were derived from dose-response curves obtained via non-linear regression analysis (see also **Figures S2B**, **C**, and **D**). The dashed diagonal line (y = x) represents values of CC50 equals to MIC. Data points in yellow indicate antimicrobial activity against *S. aureus* ATCC 12600, with point size corresponding to the log2MIC against this strain. **(E)** Ternary plots depicting the secondary structure composition (α-helix, β-sheet, and disordered) of each peptide at 50 µmol L^-1^ in different solvents: water, 60% trifluoroethanol (TFE) in water, 60% methanol (MeOH) in water, and sodium dodecyl sulfate (SDS, 10 mmol L^-1^) in water. Secondary structure fractions were estimated using the BeStSel server ^21^ (see also **Figure S3**). Figure created with BioRender.com.

Due to its ability to invade host cells, establish persistent colonization, and cause recurrent infections by replicating within both phagocytic and non-phagocytic cells ^12^, *S. aureus* was selected as an intracellular pathogen for experimental characterization in this study.

### Mechanism of action of Turingcins

To explore the antimicrobial mechanisms of the 15 Turingcins active against *S. aureus* ATCC 12600, we assessed their ability to depolarize bacterial membranes, a typical target of AMPs through non-specific lipid bilayer interactions ^13^ **(Figures 2C and S2A)**. Using the voltage-sensitive dye DiSC3-5, which fluoresces upon membrane depolarization, we identified significant perturbations in membrane potential in the case of five peptides (Turingcins 34, 35, 37, 57, and 58), representing 33% of those tested.

In contrast, nine active peptides (53%; Turingcins 2, 8, 12, 15, 17, 22, 23, 46, and 47) exhibited minimal or no significant fluorescence changes, suggesting alternative antimicrobial mechanisms. Interestingly, Turingcin-28 reduced the fluorescence of the probe, a phenomenon potentially linked to its high tryptophan content (6 out of 18 residues, 33%), the highest among the tested sequences **(Figures 2C and S2A)**.

This diversity in mechanisms of action highlights the algorithm’s ability to generate peptides with multifaceted antimicrobial strategies, reflecting a versatile approach to addressing infections.

### *In vitro* cytotoxicity and hemolytic activity

Evaluating the potential toxicities of Turingcins is a critical step in assessing their bioactivity and therapeutic potential. To assess their toxicity, we conducted dose-response 3-(4, 5-dimethylthiazolyl-2)-2,5-diphenyltetrazolium bromide (MTT) cytotoxicity assays using immortalized human keratinocytes (HaCaT) and phorbol 12-myristate 13-acetate (PMA)-differentiated human monocytes (THP-1), modeling macrophages ^14,15^. These assays assessed the biocompatibility of 43 antimicrobial Turingcins with both non-phagocytic and phagocytic cells **(Figures 2D, S2B and S2C; Table S2)**. CC50 values (the concentration causing 50% cell death) were determined through non-linear regression of dose-response data (3.12–50 μmol L^−1^) and compared to their corresponding MIC values **(Figures 2D, S2B and S2C; Table S2)**.

In HaCaT cells, 9.3% of Turingcins (4 out of 43) showed CC50 values below 12.5 μmol L^−1^, compared to only 3% (1 out of 43) in PMA-differentiated THP-1 cells, indicating greater susceptibility of keratinocytes than macrophages to Turingcins. Nearly all peptides exhibited CC50 values at least twice their MIC, except Turingcin-32, whose antimicrobial effective dose (50 μmol L^−1^) exceeded its CC50 against HaCaT cells (34.84 μmol L^−1^).

To further assess safety, we evaluated the hemolytic activity of Turingcins by exposing red blood cells (RBCs) to the peptides, a standard method for toxicity evaluation of antimicrobial agents ^15^. Four of the 43 Turingcins tested (9.3%) displayed hemolytic concentration causing 50% cell death (HC50) at values below 12.5 μmol L^−1^ **(Figures 2D and S2D; Table S2)**.

These findings highlight the promising safety profile of ApexDuo-designed compounds. Most Turingcins demonstrated CC50 or HC50 values that exceeded their MIC by 2 to over 50 times and surpassed 100 times in the case of RBCs. These findings underscore the success of ApexDuo in balancing antimicrobial potency with minimal host cell toxicity.

### Secondary structure of Turingcins

The secondary structures of short peptides are inherently dynamic, often shifting between disordered and ordered states at hydrophobic-hydrophilic interfaces. For AMPs, this structural plasticity is crucial for their interaction with biological membranes, as it underlies their functional mechanisms of action ^16^. Likewise, recent studies indicate that the secondary structural properties and conformations of CPPs profoundly influence their cellular uptake pathways and overall internalization efficiency, underscoring the importance of peptide structure in determining both antimicrobial and cell-penetrative activities ^17^.

To characterize the secondary structure of the synthesized Turingcins, Circular Dichroism (CD) spectroscopy was performed in various environments: water, trifluoroethanol (TFE) in water mixture (3:2, v:v), methanol (MeOH) in water mixture (1:1, v:v), and sodium dodecyl sulfate (SDS) in water (10 mmol L^−1^). The TFE/water and MeOH/water mixtures were used to induce distinct secondary structures: TFE usually promotes α-helical conformations by dehydrating peptide amide groups and facilitating intramolecular hydrogen bonds, while MeOH favors β-sheet conformations by interacting with the amide groups of the backbone and favoring intermolecular side chains interactions. SDS micelles were included to mimic the bilayer environment of biological membranes and provide insight into how the peptides might behave *in vivo* ^18–20^.

All Turingcins were analyzed using CD spectroscopy over a wavelength range of 260 to 190 nm **(Figures 2E and S3).** Secondary structure fractions were estimated using the BeStSel server ^21^. In water, most peptides were predominantly unstructured. Notably, however, the inactive peptides Turingcins 1, 7, and 21 showed a pronounced tendency to adopt α-helical structures (31.4%, 39%, and 29.5%, respectively). In TFE and water mixture, unstructured conformations remained dominant, whereas in MeOH and water mixture, peptides displayed enhanced β-like structural features. In the presence of SDS micelles in water, which are intended to simulate membrane-like environments, the unstructured state generally predominated, with certain peptides exceeding 65% unstructured content **(Figures 2E and S3).** In summary, Turingcins are generally unstructured and tend to β-like conformations in β-inducer medium.

### Intracellular antimicrobial activity of Turingcins

To evaluate the intracellular antimicrobial activity of the most promising Turingcins, we developed a model using HaCaT and PMA-differentiated THP-1 cells infected with intracellular *S. aureus* ATCC 12600 **(Figure S4A)**. Gentamicin was used to eliminate extracellular bacteria due to its poor cellular uptake^22–24^. To validate the model, we also tested gentamicin at 5, 100, and 500 µg mL^-1^ and confirmed that it does not eliminate intracellular *S. aureus*, even at the highest concentration tested **(Figure S4B)**.

From the MIC panel of Turingcins **(Figure 2B)**, we selected nine highly active candidates against *S. aureus*, testing them in our intracellular antimicrobial activity assays at three concentrations (12.5, 25, and 50 µmol L^-1^) **(Figures S5A-D)**. This strategy allowed us to identify time-dependent trends while excluding conditions where toxicity obscured antimicrobial efficacy. Two out of the nine Turingcins (i.e., Turingcin-46 and Turingcin- 47) tested against intracellular *S. aureus* effectively reduced bacterial loads at nontoxic concentrations, with particularly strong effects observed at 1 and 3 hours **(Figures 3A and S5E**; see also **S6A** for extracellular bacterial counts).

**Figure 3.**
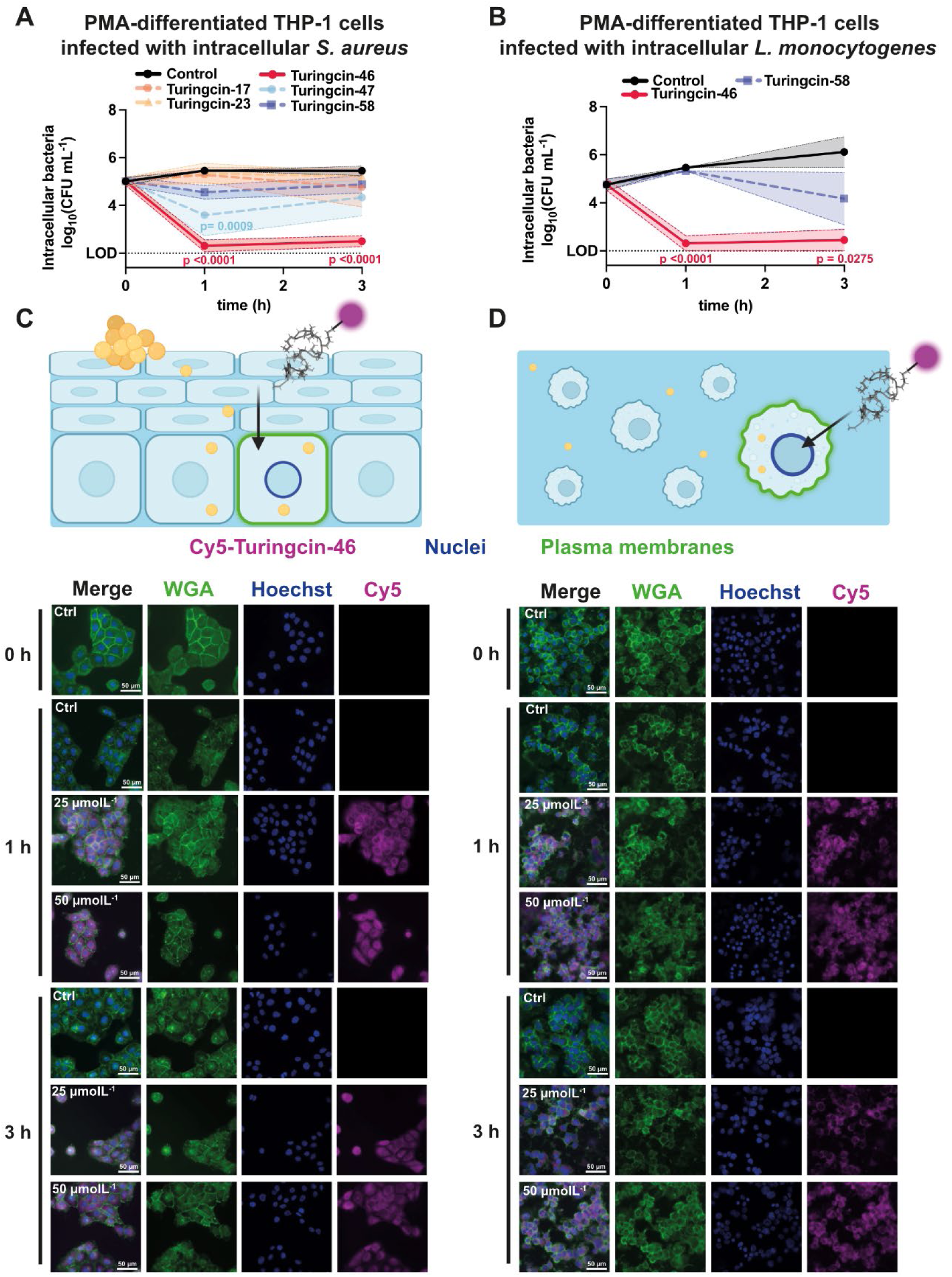
Intracellular antimicrobial activity of Turingcins. **(A**, **B)** Time-course analysis of the intracellular antimicrobial activity of Turingcin peptides at non-cytotoxic concentrations against *S. aureus* ATCC 12600 and *L. monocytogenes* ATCC 19111 in PMA-differentiated THP-1 macrophages. Intracellular bacterial loads were quantified at 0, 1, and 3 hours post-treatment (see also **Figure S6A-B** for extracellular bacteria counts). A minimum of two experiments was performed for all conditions, with the lead compound (Turingcin-46) tested in three replicates. **(C, D)** Representative fluorescence microscopy images of HaCaT and PMA-differentiated THP-1 cells treated with Cy5- labeled Turingcin-46, captured at 0-, 1-, and 3-hours post-treatment. Following peptide incubation, samples were fixed with 4% PFA and stained with WGA 488 (10 µg mL⁻¹) to visualize plasma membranes and Hoechst 33342 (1 µg mL⁻¹) for nuclei visualization. Each experiment was performed in biological duplicate, with two technical replicates each. Figures were created using BioRender.com.

To expand our studies to a highly relevant intracellular pathogen, we performed additional intracellular antimicrobial assays with *L. monocytogenes*^25,26^ using Turingcin-46 and -58, both of which were active against *L. monocytogenes* in MIC assays **(Figure 2B)** and nontoxic against THP-1 cells at the concentrations tested. In these experiments, Turingcin-46, but not Turingcin-58, significantly reduced the intracellular bacterial load compared to untreated controls, confirming its efficacy against intracellular *L. monocytogenes* **(Figures 3B**; see also **S6A** for extracellular bacterial counts).

To further examine Turingcin-46’s cell-penetration capabilities and intracellular localization, we conjugated the peptide with Cyanine-5 (Cy5) at the N-terminus and applied it to both infected and uninfected cells. Two concentrations of Cy5-Turingcin-46 (25 and 50 µmol L^-1^) were tested at 1- and 3-hours post-treatment. Afterward, cells were fixed and stained with Hoechst (blue) to label nuclei and WGA (green) to visualize cell walls, then analyzed by fluorescence microscopy. Substantial intracellular accumulation of Cy5-Turingcin-46 (purple signal) was observed in both infected **(Figures S6C and D)** and uninfected cell monolayers **(Figures 3C and D)**, compared to untreated controls. In contrast, no intracellular signal was detected when using Cy5 alone under the same experimental conditions, except for faint residual fluorescence occasionally observed in infected macrophages at 3 hours at the highest concentration tested, which may reflect nonspecific uptake through phagocytic mechanisms **(Figures S7)**.

Altogether, these findings confirm Turingcin-46’s ability to penetrate mammalian cells and effectively kill intracellular pathogens such as *S. aureus* and *L. monocytogenes*.

### Ala-scan screening of Turingcin-46 to elucidate structure-function relationships

To dissect the contributions of individual amino acids to Turingcin-46’s biological activity, we performed alanine-scan screening. Each residue in Turingcin-46 was systematically replaced with alanine, generating 18 single-substitution variants **(Figures 4, S8 and S9; Table S3)**. Alanine, with its short aliphatic side chain, was chosen to preserve peptide backbone length while minimizing disruptive interactions between side chains ^27,28^.

**Figure 4.**
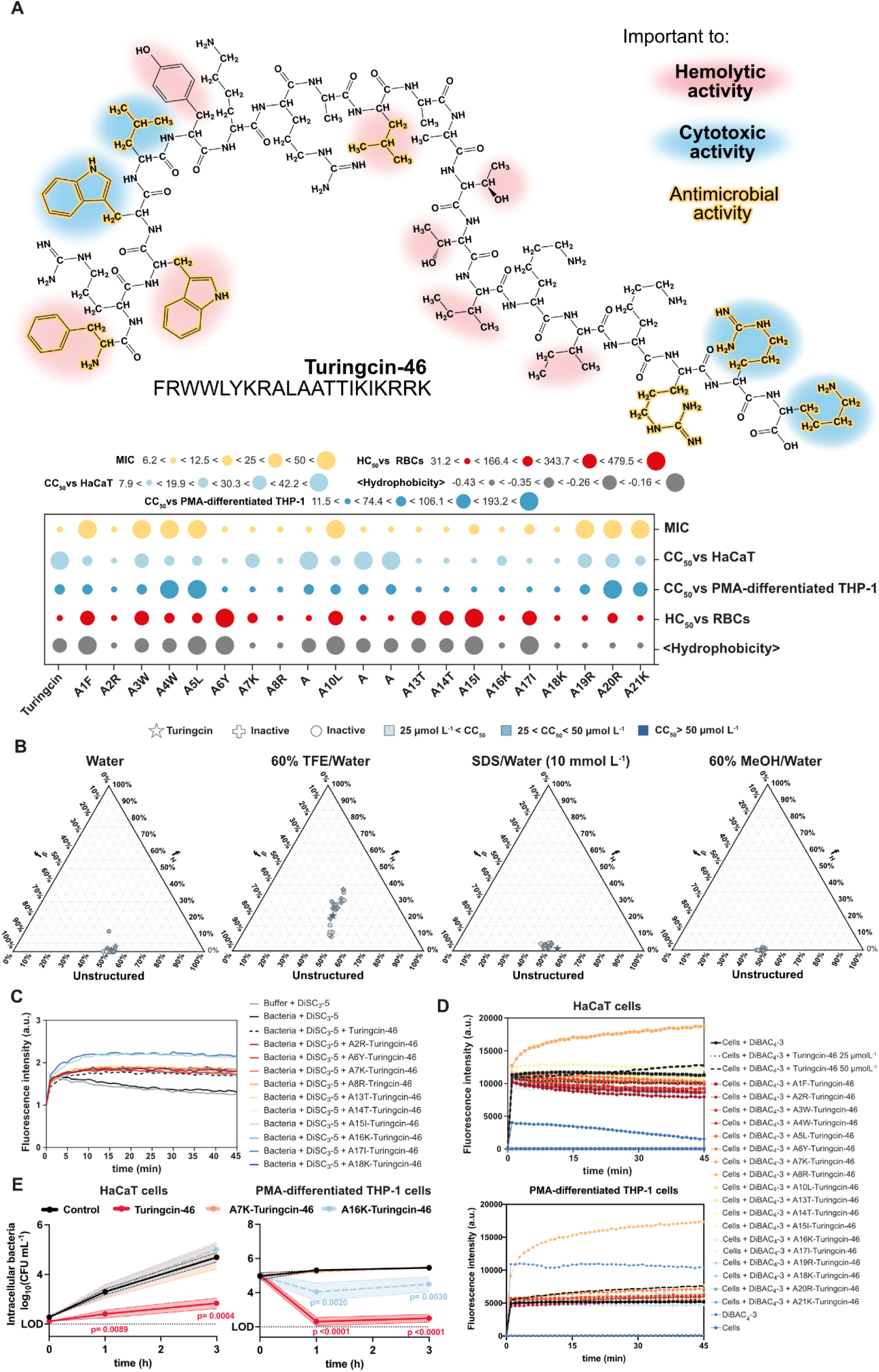
Structural and functional elucidation of Turingcin-46. **(A)** Chemical structure of Turingcin-46, with residues influencing antimicrobial activity (yellow), cytotoxicity (blue), and hemolytic effects (red) highlighted. MIC (against *S. aureus* ATCC 12600), CC50 (against HaCaT keratinocytes and PMA-differentiated THP-1 macrophages), HC50, and normalized hydrophobicity values are presented in a bubble plot to compare the wild-type Turingcin-46 sequence with 18 alanine-substituted variants (see also **Figures S8A-D**). Normalized values (ranging from 0 to 1) were used for visualization, while experimental data and normalized hydrophobicity values are provided in the legend. The peptides hydrophobicity was obtained using the DBAASP server ^39^ using the Eisenberg and Weiss scale. **(B)** Ternary plots showing the secondary structure composition of each peptide (50 µmol L^-1^) in different solvents: water, 60% trifluoroethanol (TFE) in water, 60% methanol (MeOH) in water, and sodium dodecyl sulfate (SDS, 10 mmol L^-1^) in water. Secondary structures were estimated using the BeStSel server ^21^ (see also **Figure S9A-E**). **(C)** Membrane depolarization of *S. aureus* cells treated with Turingcin-46 and alanine-substituted derivatives that retained antimicrobial activity. Changes in fluorescence intensity were measured using DiSC3-5- dye at MIC concentrations (see also **Figure S8E**). **(D)** Effects on eukaryotic cell membrane potential (depicted as depolarization or hyperpolarization) for HaCaT and PMA-differentiated THP-1 cells treated with Turingcin-46 and alanine variants, evaluated using DiBAC4-3 dye at CC50 concentrations (see also **Figures S8F-G**). **(E)** Time-course analysis (0, 1, and 3 hours) of intracellular antimicrobial activity of Turingcin-46, K7A-Turingcin-46, and K16A-Turingcin-46 at non-cytotoxic concentrations against intracellular *S. aureus* in HaCaT keratinocytes and PMA- differentiated THP-1 macrophages (see also **Figure S9F-G** for extracellular bacteria counts). All experiments were carried out in three independent replicates.

Analysis of the antimicrobial and cytotoxic properties of these alanine-substituted variants revealed that mutations at positions 1 (F), 3 (W), 4 (W), 5 (L), 10 (L), 19 (R), 20 (R), and 21 (K) significantly increased the MIC against *S. aureus* ATCC 12600, rendering the peptides inactive **(Figures 4A and S8A)**. Substitutions at positions 15 (I) and 17 (I) also diminished antimicrobial potency, albeit to a lesser degree. These findings highlight the importance of hydrophobic and aliphatic residues such as tryptophan, phenylalanine, leucine, and isoleucine, as well as positively charged arginine and lysine, in mediating antimicrobial effects. Notably, the changes in cationic residues close to the C-terminus were more impactful than close to the N-terminus where cationic residues are surrounded by hydrophobic and aliphatic residues.

Some of the residues affecting MIC values also impacted cytotoxicity **(Figures 4A, S8B and S8C)**. Mutations leading to a loss of antimicrobial activity generally reduced cytotoxicity in PMA-differentiated THP-1 cells (e.g., W4A, L5A, R20A, and K21A). All the derivatives had similar activity profiles or were more toxic than Turingcin-46 against keratinocytes. The wild-type peptide did not present high hemolytic activity, and the Ala- scan screening showed that substitutions that led to enhanced mean hydrophobicity values correlated with a lower propensity for hemolysis **(Figures 4A and S8D; table S4)**.

Circular dichroism analysis showed that Turingcin-46 and its alanine-substituted variants generally adopted similar secondary structures **(Figures 4B and S9)**. In aqueous solution, the peptides were predominantly unstructured, with some β-like conformations. In TFE and water mixture, variants that lost antimicrobial activity showed an increased tendency to form α-helical structures (up to 37%). Nevertheless, in MeOH and water mixture and SDS micelles, both wild-type and mutated sequences displayed similar proportions of unstructured and β-like conformations **(Figures 4B and S9A-E)**.

To examine how these structural changes affected interactions with biological membranes, we employed two fluorescent probes—DiSC3-5 and DiBAC4-3 ^29,30^—to evaluate membrane depolarization in bacterial and eukaryotic cells, respectively. Turingcin-46 had a modest ability to depolarize the bacterial membrane, a trait shared by most alanine-substituted variants **(Figures 4C and S8E)**. Interestingly, I15A- and I17A- substituted Turingcin-46 displayed enhanced depolarizing activity against *S. aureus* membranes **(Figures 4C and S8E)**.

The peptides exhibited distinct effects on eukaryotic cell membranes. In HaCaT keratinocytes, Turingcin-46 did not alter membrane potential, whereas most alanine-substituted derivatives induced hyperpolarization (indicated by decreased fluorescence), particularly those mutated at the N- and C-termini. Conversely, K7A- and I15A- substituted variants increased fluorescence, indicating membrane depolarization. In PMA-differentiated THP-1 macrophages, wild-type Turingcin-46 induced membrane depolarization, whereas most alanine-substituted variants showed reduced depolarizing effects **(Figures 4D and S8F-G)**. One exception was K7A-Turingcin-46, which caused significantly greater depolarization than the wild-type sequence. These results underscore the cell-type specificity of peptide-membrane interactions influenced by phospholipid composition.

To further assess intracellular antimicrobial activity, we selected the K7A- and K16A- Turingcin-46 variants based on their antimicrobial potency and cytotoxicity profiles, which were comparable to those of the lead peptides. However, K7A-Turingcin-46 did not reduce the intracellular bacterial burden in HaCaT and THP-1 cells when tested at non-cytotoxic concentrations **(Figure 4E**; see also **Figure S9F-G** for extracellular bacteria counts). Conversely, K16A-Turingcin-46 preserved intracellular activity in macrophages, although its effectiveness was diminished compared to the lead wild-type sequence **(Figure 4E**; see also **Figure S9F-G** for extracellular bacteria counts). This suggests that these variants lack key sequence features necessary for the full multimodal activity of the wild-type peptide.

In summary, alanine-scan screening revealed the importance of Turingcin-46’s unique combination and distribution of amino acid residues in mediating its membrane interactions and multifunctional biological activity. This analysis highlights the nuanced interplay between sequence composition and peptide function in computationally designed molecules.

### Resistance to proteolytic degradation

Before conducting *in vivo* mouse experiments, we assessed Turingcin-46’s stability in the presence of serum proteases. After one hour of incubation, approximately 50% of the peptide remained intact. We attribute this stability to the presence of hindered hydrophobic residues such as phenylalanine and tryptophan that may further reduce protease accessibility **(Figure S10A).**

### Anti-infective efficacy of Turingcin-46 in animal models

To investigate Turingcin-46’s antimicrobial efficacy in complex biological environments, we tested it in two mouse models: a skin abscess model and a peritonitis/sepsis model **(Figures 5, S10B-D** and **S11)**.

**Figure 5.**
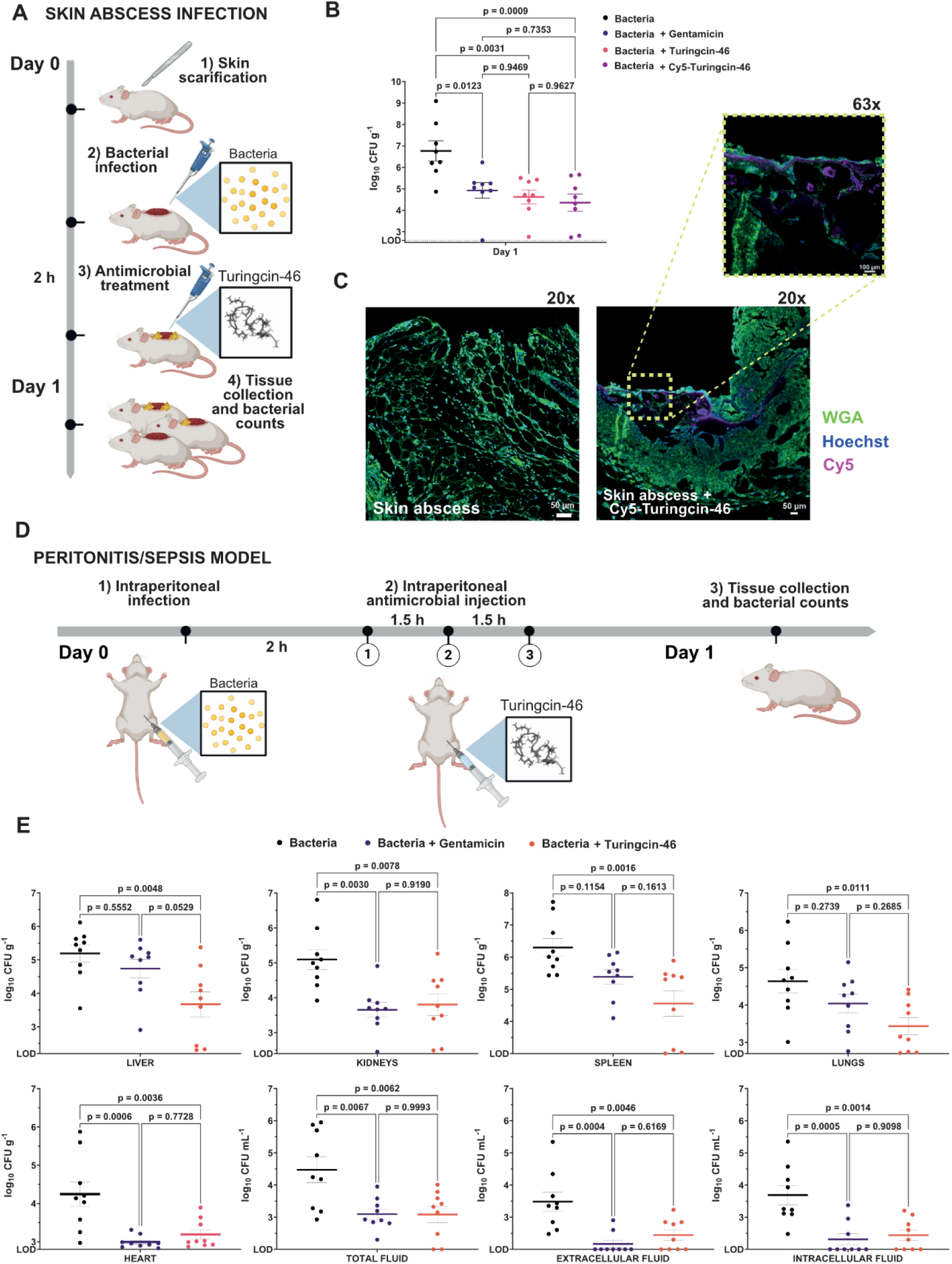
Anti-infective activity of Turingcin-46 in animal models. **(A)** Schematic representation of the skin abscess mouse model used to evaluate the anti-infective efficacy of Turingcin-46 against *S. aureus* ATCC 12600. **(B)** In the skin abscess model, a single post-infection administration of Turingcin-46 significantly reduced bacterial load compared to the untreated control. **(C)** Representative confocal laser scanning microscopy (CLSM) images of excised skin tissues from untreated mice and those treated with Cy5-Turingcin-46. Deparaffinized tissue sections were stained with WGA 488 (10 µg mL^-1^) to visualize plasma membranes and Hoechst 33342 (1 µg mL^-1^) for nuclei. **(D)** Schematic representation of the peritonitis/sepsis mouse model, in which Turingcin-46 was administered intraperitoneally. **(E)** Anti-infective efficacy against *S. aureus* ATCC 12600 was evaluated one day after injection of Turingcin-46, demonstrating significant efficacy in in multiple organs (liver, kidneys, spleen, lungs, heart) and peritoneal fluids (total, extracellular, intracellular - see also **Figure S11** for correlations between intracellular and extracellular *S. aureus* CFU counts in peritoneal lavage samples). Compared to the untreated control, Turingcin-46 treatment significantly decreased bacterial loads. Statistical significance in **(B, E)** was determined using one-way ANOVA followed by Tukey’s test; p values are shown in the graphs. The solid line within each distribution represents the mean. Panels **(A)** and **(D)** were created with BioRender.com.

In the skin abscess model, a 20 μL suspension containing 10^7^ *S. aureus* ATCC 12600 cells in phosphate-buffered saline (PBS) was applied to a wound on the back of the mice ^11,31–33^ **(Figures 5A, 5B and S10B)**. A single dose of Turingcin-46 was administered directly to the infected area. After 24 hours, the bacterial count decreased by more than two orders of magnitude compared to the untreated control group, with Turingcin-46 demonstrating slightly greater potency than gentamicin **(Figure 5B)**. Confocal microscopy of skin samples treated with Cy5-labeled Turingcin-46 confirmed the peptide’s cellular penetration capability, showing fluorescence within the stratified squamous epithelium **(Figures 5C and S10C)**. Throughout the experiments, no significant changes in body weight, a general indicator of potential toxicity, were detected **(Figure S10B)**.

Next, we evaluated Turingcin-46 in a peritonitis/sepsis mouse model **(Figures 5D, 5E, S10D** and **S11A)**, which allows for the evaluation of intracellular antimicrobial activity ^34,35^. Mice were intraperitoneally infected with 100 μL containing 5×10^10^ *S. aureus* ATCC 12600 cells *per* mL. Two hours post-infection, three doses of Turingcin-46 were administered intraperitoneally at 1.5-hour intervals to mitigate peptide degradation (as confirmed by prior proteolytic stability studies).

One day after treatment, Turingcin-46 significantly reduced bacterial burdens in multiple organs, including the liver, kidneys, spleen, lungs, and heart, with reductions ranging from 1.2 to 1.8 orders of magnitude. Notably, Turingcin-46 outperformed gentamicin (positive control) in the liver, spleen, and lungs, suggesting possible organ-specific peptide accumulation. Importantly, no significant changes in body weight were observed, further supporting the peptide’s safety at the tested dose **(Figure S10D)**.

Gentamicin has poor uptake into mammalian cells, as confirmed by our *in vitro* data **(Figure S4B)** and supported by numerous previous studies ^22–24^, making it unlikely to act directly on intracellular bacteria. However, varying degrees of positive correlation were observed between intracellular and extracellular bacterial loads across treatment groups *in vivo* **(Figure S11B)**, suggesting that the extracellular burden may influence intracellular load through reinfection. A linear regression analysis revealed that the slope of intracellular versus extracellular bacterial loads was substantially lower in the Turingcin-46 group (0.85) compared to the control (1.03) and gentamicin (1.37) groups, indicating a disproportionately stronger reduction in intracellular bacteria counts in the Turingcin-46 treatment group. In contrast, the steeper slope observed in gentamicin-treated mice suggests that its apparent effect on intracellular bacteria may primarily result from the reduction of extracellular burden. Moreover, the intracellular-to-extracellular bacterial load ratio in the Turingcin-46 group was 1.74-fold lower than that of the gentamicin group, further supporting Turingcin-46’s intracellular antimicrobial activity through this comparative trend **(Figure S11C)**.

Overall, these *in vivo* findings highlight Turingcin-46’s potent antibiotic activity under physiological conditions and highlight its potential as a multimodal therapeutic agent against infections.

## Discussion

Eliminating intracellular pathogens remains a significant therapeutic challenge. Traditional antibiotics often fail to reach sufficient concentrations within host cells to effectively eradicate intracellular bacteria, and achieving therapeutic intracellular levels frequently requires high, potentially toxic doses ^36^.

While multimodal models are now widely applied in domains such as chatbots, their use in molecular design has been limited due to the complexity of integrating multiple functionalities into a single molecule. Multimodal molecules offer a promising strategy to overcome these barriers, but their design and optimization using conventional methods have proven difficult.

ApexDuo introduces a dual-objective deep learning framework that generates peptides with both antimicrobial and cell-penetrating activities. Whereas most molecular discovery pipelines optimize a single property, ApexDuo couples predictive models for multiple bioactivities within a unified generative workflow, enabling the rational design of functionally versatile peptides. Although antimicrobial and cell-penetrating properties are often studied separately, they are not mutually exclusive ^37,38^. ApexDuo exploits this overlap by deliberately designing peptides that combine both functionalities. The Turingcin series discovered through this approach displays a hybrid chemical profile characteristic of antimicrobial and cell-penetrating peptides, demonstrating the efficacy of the strategy. Looking ahead, the framework can be expanded to incorporate further biological functions, such as immunomodulation, endosomal escape, or subcellular targeting, thereby supporting the programmable design of next-generation therapeutics for complex infectious and inflammatory diseases.

Mutational analysis identified sequence elements critical for Turingcin-46’s antimicrobial efficacy and cytotoxic selectivity, correlating these features with the peptide’s ability to penetrate mammalian cells, suppress intracellular pathogens, and maintain a favorable cytotoxic profile. Furthermore, *in vivo* experiments demonstrated Turingcin-46’s robust antimicrobial efficacy in two mouse models, underscoring its potential therapeutic potential. In summary, ApexDuo demonstrates that AI can be used to design multimodal peptides, opening new avenues for designing molecules with more than one function.

## Limitations of the study

Despite the advances outlined here, ApexDuo has limitations. The antimicrobial predictor did not incorporate datasets specifically annotated for intracellular activity, which may have constrained the accuracy of predictions for this context. Similarly, limited publicly available data on CPPs restricted the predictor’s training. Future efforts to curate large datasets with intracellularly active peptide and cell-penetration annotations will enhance predictive models and accelerate the discovery of improved multimodal antibiotics. Additionally, our study focused on peptides with confirmed antimicrobial activity for downstream intracellular assays, excluding purely cell-penetrating candidates. Broader testing could provide further insights into the full spectrum of ApexDuo-generated molecules and identify additional promising candidates. Finally, we provide strong evidence that lead candidates (i.e., Turingcin-46), significantly reduce infections in mammalian cells and in murine models. Future studies should more granularly delineate this compound’s intracellular targeting across additional mouse models.

## Acknowledgments

Research reported in this publication was supported by the NIH R35GM138201 and DTRA HDTRA1-21-1-0014. We thank Dr. Mark Goulian for kindly donating the following strains: *Escherichia coli* AIC221 (*Escherichia coli* MG1655 phnE_2::FRT [control strain for AIC222]) and *Escherichia coli* AIC222 (*Escherichia coli* MG1655 pmrA53 phnE_2::FRT [polymyxin resistant]), and the H-MARC core in the Center for Molecular Studies in Digestive and Liver Diseases (P30 DK050306) for kindly donating strains *Acinetobacter baumannii* ATCC 19606 and *Klebsiella pneumoniae* ATCC 13883. We would like to acknowledge the Center for Molecular Studies in Digestive and Liver Diseases (P30DK050306) and the Molecular Pathology and Imaging Core (RRID: SCR_022420) for their assistance with histology sample preparation. Additionally, we thank the Cell & Developmental Biology Microscopy Core at the Perelman School of Medicine (RRID: SCR_022373) for enabling the use of the Zeiss LSM 980 confocal microscope. We thank Gui-Shuang Ying from the Center for Preventive Ophthalmology and Biostatistics (CPOB) Consulting Service for advice on our statistical analyses. We thank the de la Fuente lab members for insightful discussions, especially Sufen Li, for her assistance with animal experiments. Figures created with Biorender.com are attributed as such. Molecules were rendered using the PyMOL Molecular Graphics System, Version 3.2 Schrödinger, LLC.

## Author contributions

Conceptualization: AC, CFN

Methodology: AC, FW, MDTT, CFN

Experimental investigation: AC, MDTT

Computational investigation: FW

Visualization: AC, FW, MDTT

Funding acquisition: CFN

Supervision: CFN

Formal analysis: AC, FW, MDTT

Writing – original draft: AC, FW, MDTT, CFN

Writing – review & editing: AC, FW, MDTT, CFN

## Data and code availability

All public databases used for building ApexDuo are listed in the key resources table. Pretrained models for ApexDuo are available on Gitlab (https://gitlab.com/machine-biology-group-public/apex_duo) and listed in the key resources table. Any additional information needed to reanalyze the data presented in this paper can be obtained from the lead contact upon request.

## Methods

### Datasets

Peptides used for training the deep generative model were curated from UniRef50 ^40^. We filtered out protein sequences having more than 50 amino acids. The peptides we downloaded from UniRef50 were divided into a training set and a validation set with 1,418,011 and 354,503 peptides, respectively. AMPs used for updating antimicrobial predictor APEX consists of two parts. One part is the DBAASP ^39^ dataset used in the original APEX work ^49^, which contained 14,738 measured antimicrobial activities (MICs) values for 988 peptides against 34 bacterial strains. The other part is our updated in-house antimicrobial activity dataset with 1,016 peptides, 37 strains (three new strains including *Porphyromoras gingivalis* ATCC BAA-308, *Neisseria meningitis* ATCC 13077, and *E. coli* K1 ATCC 700973) and 14,963 MIC values. Public AMPs used for computational analysis were retreived from public databases, including APD3 ^41^, DBAASP ^42^ and DRAMP ^43^, as well as our in-house peptides having minimum inhibitory concentration ≤128 μmol L^-1^ against at least one bacterial strain ^49^. CPPs used for training and computational analysis were retreived from CPPsite 2.0 ^44,50^, SkipCPP-Pred ^45^ and MLCPP ^46^. We also curated toxic peptides from SATPdb ^47^ and labeled them as negative data so that the CPP predictor can balance the cell-penetrating property and toxicity and help to retrieve peptides with good cell penetration and relatively low toxicity. After cleaning the non-canonical peptides and removing duplicates, we obtained 1,217 CPPs and 5,293 non-CPPs or toxic peptides for CPP predictor building.

### Variational autoencoder

We use a maximum mean discrepancy variational autoencoder (MMD-VAE) as our peptide sequence generator ^51^. The MMD-VAE consists of an encoder *E*(⋅) and a decoder *D*(⋅), both typically implemented as neural networks. The encoder and decoder form a compression and de-compression process that enables representation learning, distribution learning and novel data generation. In our scenario, the inputs to the encoder are peptide sequences. For a peptide sequence with *l* amino acids, two special characters indicating the start and end of the sequence are added at the beginning and end, respectively. One-hot encoding is then used to numerically represent this sequence. Let *x* denotes this representation, the encoder takes *x* and generates a continuous vector (i.e., a latent code) *v* = *E*(*x*) ∈ ℝ^*d*^, where *d* stands for the dimension of *v*. The decoder uses the latent code *v* to generate a peptide sample *x*^′^ = *D*(*v*) that resembles *x*. Given a peptide set to train the MMD-VAE and let 𝑝*_data_*(*x*) denote the empirical distribution of training peptides, the minimization goal of MMD-VAE can be written as

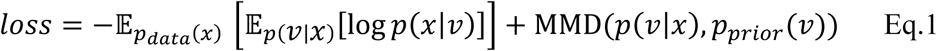

where 𝔼[⋅] stands expectation, *p*(*v*|*x*) is the posterior distribution of the latent codes given *x*𝑙 from 𝑝_*d*_(*x*), MMD is the maximum mean discrepancy that measures the distance of two probability distributions, and 𝑝*_prior_*(*v*) is the prior distribution of *v*. In our case, the log-likelihood log 𝑝(*x*|*v*) in Eq.1 is equal to the negative cross entropy between *x* from 𝑝_*d*ata_(*x*) and its reconstructed version by *D*(*E*(*x*)) the MMD-VAE. In this sense, Eq.1 can be re-written as

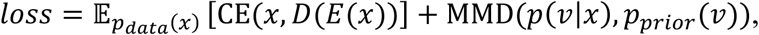

where CE(⋅,⋅) is the cross entropy. Optimizing the parameters of *E*(⋅) and *D*(⋅) to minimize 𝔼*_data_*_(*x*)_ [CE(*x*, *D*(*E*(*x*))] ensures the MMD-VAE to capture the distribution of training peptide sequences (hence, the decoder can generate peptides resemble the training ones) while the optimization of *E*(⋅) and *D*(⋅) for MMD(𝑝(*v*|*x*), 𝑝*_data_*(*v*)) makes the latent codes *v*s close to the prior distribution (hence, making it easier for the users to sample latent codes from the prior distribution and generate new peptide samples). In our practice, the prior distribution is the zero-mean, unit-variance multivariate Gaussian distribution. The encoder begins by mapping the one-hot encoding of a peptide to a learnable embedding layer 𝒙 ∈ ℝ^𝐿𝐿×𝑒𝑒^, where 𝐿 stands for the input length and 𝑒𝑒 denotes the feature dimension. Then the encoder uses a recurrent neural network (implemented as gated recurrent units) to map *x* to hidden features 𝒉 ∈ ℝ^𝐿×𝑘^, where 𝑘𝑘 denotes the number of hidden units. We then adopt an attention mechanism to fuse contextual residue information by

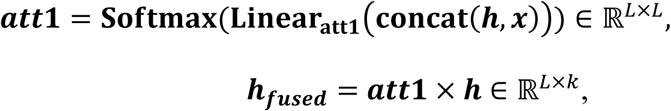

where **Linear_att1_**(⋅) is a linear transformation, **concat**(⋅,⋅) is concatenation operation along the feature dimension (i.e., the last dimension), **Softmax**(⋅) is the softmax function and also applied to the last dimension of its input. After that, we use another attention mechanism to final latent code 𝒉*_latent_* of the peptide by

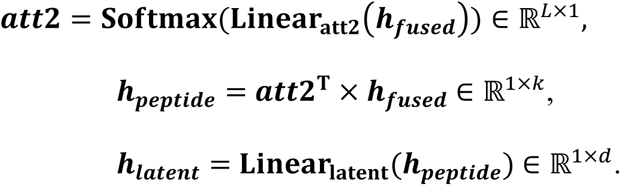

The decoder is a recurrent neural network with the same architecture as that from the encoder followed by a fully-connected neural network layer to transform the latent code to the residue type probabilities per amino acid position. We used a learning rate of 0.0001 to train the MMD-VAE 1,000 epochs on the training set. The number of recurrent neural network layers, the number of hidden units in the recurrent neural network, the dimension of the latent code, batch size, weight decay strength, and the best epoch were chosen by the *llll* of the trained model evaluated on the validation set.

### Peptide generation

Instead of directly sampling latent codes to generate peptide samples, we followed the previous practice to fit a kernel density estimator (KDE) to the functional peptides. That is, we obtained the latent codes of our in-house peptides and publicly available CPPs. A KDE (implemented by scikit-learn with default hyperparameter) was used to fit the distribution of these latent codes. We then sampled the latent codes from the KDE and decoded the sampled codes by the decoder to obtain the generated peptide sequences. We sampled 50M latent codes and generated the corresponding sequences for further filtering by the antimicrobial and cell-penetrating peptide predictors.

### Antimicrobial peptide predictor

The antimicrobial peptide predictor we used is APEX ^49^, a multi-bacterial multi-strain antimicrobial peptide predictor previously been built for mining antibiotic encrypted peptides from proteomes of extinct organism. APEX uses the same architecture of the encoder of our variational autoencoder except we drop the embedding layer and use AAindex^52^ descriptors of amino acid sequences as inputs. That is, given an input peptide, it is represented as 𝒙 ∈ ℝ^𝐿𝐿×566^, where 𝐿 is peptide length and ith column of 𝒙 is a 566- dimensional vector storing the physicochemical and biochemical properties of ith amino acid in the peptide. After extracting the 𝒉*_latent_*, we apply multilayer perceptron to process 𝒉*_latent_* and predict antimicrobial peptides. For each generated peptide, we used APEX to obtain the median value of predicted minimum inhibitory concentrations (MICs) against multiple bacterial strains.

### Cell-penetrating peptide predictor

We used deep learning to build the binary classifier to predict the cell-penetrating property of peptide sequences. Our predictor has similar architecture to that of the encoder of MMD-VAE except that we added one more fully-connected neural network layer to the hidden peptide features to the cell-penetrating probabilities. We set the learning rate = 0. 0001, batch size = 128, the number of training epochs = 100, and trained each model five times under different seeds to create an ensemble version of it (i.e., we averaged the prediction result from each model). Hyperparameters (i.e., the number of recurrent neural network layers, the hidden unit size, and weight decay strength) of the CPP predictor were selected based on the mean of the area under the receiver operating characteristic under the five-fold cross validation. The CPP predictors from the top five hyperparameter combinations were selected (i.e., 25 models = 5 hyperparameter combinations × 5 copies under different random seeds) for the CPP peptide screening.

### Peptide selection

We used the following rules to filter and select the desired peptides from 50M generated peptides from the MMD-VAE.

1. Generated peptides having median MIC >32 ìmol L^-1^ were filtered out from our selection.
2. After screening the CPP probabilities for the 50M generated peptides, we calculated the mean and standard deviation of the predicted probabilities among the 50M sequences. Generated peptides passing 3-sigma rule (i.e., CPP prediction ≥ mean of CPP prediction + 3 × standard deviation of CPP prediction) were kept.
3. Generated peptides having more than 30 amino acid residues were filtered out due to synthesis difficulty.
4. Generated peptides appeared in our in-house peptide data or training CPPs were removed.
5. Remaining generated peptides were ranked by median MIC, and we used Striped Smith–Waterman algorithm to perform pair-wise sequence alignment and calculate the sequence similarity between two peptides. We kept the peptide with lower median MIC when a pair of peptides having a sequence similarity > 0.5.

### Physicochemical properties extraction

The twelve physicochemical properties of the peptides, including normalized hydrophobic moment, normalized hydrophobicity, net charge, amphiphilicity index, isoelectric point, penetration depth, tilt angle, disordered conformation propensity, linear moment, tendency for in vitro aggregation, angle between hydrophobic residues, and propensity for PPII coil, were obtained using the DBAASP server ^39^. The hydrophobicity scale used was the Eisenberg and Weiss scale.

### Peptide synthesis

The peptides used in the experiments were procured from AAPPTec and produced through solid-phase synthesis employing the Fmoc strategy.

### Bacterial strains and growth conditions

In this study, we worked with the following pathogenic bacterial strains: *Staphylococcus aureus* ATCC 12600, *Staphylococcus aureus* ATCC BAA-1556 (methicillin-resistant strain), *Enterococcus faecalis* ATCC 700802 (vancomycin-resistant strain), *Enterococcus faecium* ATCC 700221 (vancomycin-resistant strain), *Listeria monocytogenes* ATCC 19111, *Salmonella enterica* ATCC 9150, *Escherichia coli* ATCC 11775, *Escherichia coli* AIC221 [*Escherichia coli* MG1655 phnE_2::FRT (control strain for AIC222)], *Escherichia coli* AIC222 [*Escherichia coli* MG1655 pmrA53 phnE_2::FRT (polymyxin resistant; colistin-resistant strain)], *Acinetobacter baumannii* ATCC 19606, *Klebsiella pneumoniae* ATCC 13883, and *Pseudomonas aeruginosa* PAO1. All strains were cultured in Luria-Bertani (LB) broth or on LB agar, except for *L. monocytogenes*, which was cultured in Brain Heart Infusion (BHI) medium, and *S. aureus*, which was grown on mannitol salt agar plates. Bacterial cultures for all experiments were started from a single isolated colony and grown overnight (16 hours) in liquid medium at 37 °C. The next day, the cultures were diluted 1:100 into fresh medium and incubated at 37 °C until reaching the mid-logarithmic growth phase.

### Minimal inhibitory concentration assays

Broth microdilution assays were conducted to determine the MIC, the minimum inhibitory concentration required to achieve complete bacterial elimination, for each peptide. Peptides were prepared in sterile water and serially diluted twofold in untreated polystyrene 96-well plates, with final concentrations ranging from 0 to 100 µmol L^-1^, as previously described^11,53^. Bacterial cultures, adjusted to 4 × 10^6^ CFU mL^-1^ in LB or BHI medium, were combined with the peptide solutions in equal volumes. After 24 hours of incubation at 37 °C, absorbance at 600 nm was measured using a spectrophotometer. The MIC was identified as the lowest peptide concentration that prevented visible bacterial growth. Negative control wells without peptide were included to confirm uninhibited bacterial growth.

### Bacterial membrane depolarization assays using DiSC_3_-5

A cytoplasmic membrane depolarization assay was carried out using the membrane potential-sensitive dye 3,3’-dipropylthiadicarbocyanine iodide (DiSC3-5, Sigma 43608). Mid-logarithmic phase *S. aureus* ATCC 12600 was washed and adjusted to an optical density of 0.05 at 600 nm in HEPES buffer (pH 7.2) supplemented with 20 mmol L⁻¹ glucose and 0.1 mol L^-1^ KCl. The bacterial suspension (100 µL per well) was incubated with 10 µmol L^-1^ DiSC3-5 for 15 minutes. This allowed fluorescence to stabilize as the dye incorporated into the bacterial membrane. The peptides were subsequently added at concentrations corresponding to their MIC values in a 1:1 ratio with the bacterial suspension. Fluorescence changes (λₑₓ = 622 nm, λₑₘ = 670 nm) were recorded over 45 minutes to monitor membrane depolarization.

### Eukaryotic cell culture conditions

Immortalized human keratinocytes (HaCaT, sourced from Deutsches Krebsforschungszentrum - DKFZ ^48^) and human monocytes (THP-1, ATCC TIB-202) were maintained in high-glucose Dulbecco’s modified Eagle’s medium (DMEM) and Roswell Park Memorial Institute 1640 (RPMI), respectively. Both media were enriched with 1% antibiotics (penicillin/streptomycin) and 10% fetal bovine serum (FBS). The cells were cultured at 37 °C in a humidified incubator with 5% CO2.

### Cytotoxicity assays

For HaCaT cells, 5 × 10^3^ cells per well were seeded into 96-well plates one day prior to treatment with increasing peptide concentrations (3.12–50 μmol L^-1^). THP-1 monocytes were differentiated into macrophages by treating them with 100 nmol L^-1^ phorbol 12- myristate 13-acetate (PMA) for three days before administering the peptides. After a 24- hour incubation with each peptide, an MTT assay was performed. Briefly, MTT reagent (0.5 mg mL^-1^) in phenol red-free medium (DMEM for HaCaT cells and RPMI for PMA-differentiated THP-1 cells) was added to each well (100 μL per well). The plates were incubated at 37 °C for 4 hours to facilitate the formation of insoluble formazan salts. These salts were subsequently dissolved in a solution of 0.04 mol L^-1^ hydrochloric acid (HCl) in anhydrous isopropanol, and the absorbance at 570 nm was recorded using a spectrophotometer.

### Hemolysis assays

To assess the release of hemoglobin from human red blood cells (RBCs) following treatment with each peptide, human erythrocytes (blood type A-, obtained from ZenBio, using heparin-anticoagulated blood) were washed three times with PBS (pH 7.4) by centrifugation at 800 g for 10 minutes. A 1:200 dilution of RBCs (75 μL) was combined with different concentrations of peptide solution (0.78–100 μmol L^-1^; 75 μL), and the resulting mixture was incubated at room temperature for 4 hours. After incubation, the mixture was centrifuged at 1,300 g for 10 minutes to pellet the cells and debris. The supernatant (100 μL) from each well was moved to a new 96-well plate for absorbance analysis at 405 nm using an automated plate reader. The percentage of hemolysis was calculated by comparing the absorbance to the negative control (PBS) and the positive control (1% SDS in PBS, v/v).

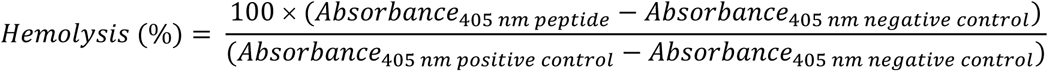

### Circular dichroism experiments

Circular dichroism (CD) measurements were performed using a J1500 CD spectropolarimeter (Jasco) at the Biological Chemistry Resource Center (BCRC) of the University of Pennsylvania. The experiments were carried out at 25 °C, and the spectra shown are the result of averaging three accumulations. A quartz cuvette with a 1.0 mm optical path length was used, covering a wavelength range of 260–190 nm at a scanning speed of 50 nm min^-1^ and a bandwidth of 0.5 nm. Peptide samples were tested at a concentration of 50 µmol L^-1^ in various environments: water, a 60% trifluoroethanol (TFE) in water mixture, a 50% methanol (MeOH) in water mixture, and water containing sodium dodecyl sulfate (SDS) at 10 mmol L^-1^. Baseline readings were taken before data acquisition, and a Fourier transform filter was used to minimize background noise. The secondary structure fraction values were calculated using the single-spectrum analysis tool on the BeStSel server ^21^. Ternary plots were created in https://www.ternaryplot.com/ and subsequently edited.

### Intracellular antimicrobial activity

HaCaT and PMA-differentiated THP-1 cells were seeded at a density of 50,000 cells per well (100 µL) in antibiotic-free medium. Cells were incubated for 2 hours with *S. aureus* ATCC 12600 or *L. monocytogenes* to allow infection, using initial bacterial densities of 0.002 or 0.01 OD600 mL^-1^ for keratinocytes, and 0.001 or 0.002 OD600 mL^-1^ for macrophages, respectively. After the incubation period, gentamicin was added at a final concentration of 500 µg mL^-1^ to remove extracellular bacteria. Following gentamicin removal, samples were treated with peptides at concentrations of 12.5, 25, and 50 µmol mL^-1^ or with gentamicin (as a negative control) at 5, 100, and 500 µg mL^-1^. At designated time points (0, 1, and 3 hours post-treatment), the supernatants were collected, serially diluted 10-fold, and plated on LB agar for colony counting. The cellular pellets were washed with PBS, lysed using 1% Triton X-100, and similarly plated on LB agar for bacterial enumeration.

### Fluorescence microscopy analysis of Cy5-Turingcin-46 internalization in eukaryotic Cells

HaCaT and PMA-differentiated THP-1 cells were seeded on glass coverslips in 24-well plates at densities of 50,000 and 400,000 cells per well, respectively. Experimental conditions were designed to evaluate Cy5-Turingcin-46 internalization in both infected and uninfected cells. For the infection condition, cells were incubated for 2 hours with *S. aureus* ATCC 12600 at an initial bacterial density of 0.02 OD mL^-1^ to enhance bacterial visualization under the microscope. Following bacterial exposure, cells were treated with gentamicin for an additional 2 hours as previously described. After treatment, cells were washed with PBS and incubated with Cy5-Turingcin-46 or Cy5 alone at concentrations of 25 and 50 µmol L^-1^. In parallel, In parallel, treatments with peptide or Cy5 alone were performed on uninfected cells that were not exposed to *S. aureus*. At three time points (0, 1, and 3 hours post-treatment), cells were washed with PBS and fixed with 4% paraformaldehyde in PBS for 15 minutes at room temperature. Nuclei, cell walls, and bacteria were stained using Hoechst 33342 (1 µg mL^-1^) and wheat germ agglutinin conjugated to Alexa Fluor 488 (WGA 488, 10 µg mL^-1^) for 20 minutes at room temperature. The stained cells were washed twice with PBS, mounted on glass slides, and imaged using a Nikon fluorescence microscope.

### Eukaryotic membrane depolarization assays using DiBAC_4_-3

HaCaT and PMA-differentiated THP-1 cells were seeded at a density of 50,000 cells per well in 96-well plates (Corning 3904) using phenol red-free medium to minimize background fluorescence. Cellular monolayers were washed twice with 100 µL of a physiological buffer composed of 140 mmol L^-1^ NaCl, 5.4 mmol L^-1^ KCl, 1 mmol L^-1^ CaCl2, 1 mmol L^-1^ MgCl2, 10 mmol L^-1^ glucose, and 10 mmolL^-1^ HEPES (pH 7.4) to remove residual medium. The cells were subsequently incubated with 5 µmol L^-1^ DiBAC4-3, a voltage-sensitive anionic dye (Thermo Fisher, catalog no. B438), prepared in the same buffer. This incubation was conducted at 37°C for 40 minutes to ensure adequate dye uptake and equilibration. Following incubation, the monolayers were gently washed to remove excess dye before being exposed to peptides at their CC50 concentrations. Fluorescence changes (λₑₓ = 490 nm, λₑₘ = 516 nm) were recorded in real-time for 45 minutes using a microplate reader, providing a quantitative assessment of membrane depolarization dynamics.

### Resistance to proteolytic degradation assay

The stability of Turingcin-46 against enzymatic degradation was assessed by incubation in a solution containing 25% human serum in water. Specifically, the peptide, at a concentration of 7 mg mL^-1^, was incubated with an aqueous solution of 25% human serum (Zen-Bio; healthy donor, blood type A-) at 37 °C for 6 hours. Samples were collected at 0.5, 1, 2, 4, and 6 hours, and each aliquot was immediately treated with 10 μL of trifluoroacetic acid in ice for 10 minutes to halt enzymatic activity. Peptide analysis was performed using a Waters Acquity ultra-high-performance liquid chromatography-mass spectrometry (UHPLC-MS) system featuring a photodiode array detector (190–400 nm data collection) and a Waters single quadrupole detector 2 (SQD2). The setup included a Waters XBridge C18 column (3.5 µm, 4.6 mm x 50 mm), with a mobile phase consisting of 100% water containing 0.1% (v/v) formic acid (solvent A) and 100% acetonitrile (solvent B). Both solvents were of Fisher Optima grade. A 20 μL injection volume was used, and ionization was carried out in both positive and negative ESI modes, scanning a mass range of m/z 100–3,000. The gradient used consisted in 5-95% solvent B in 6 min. The proportion of intact peptide remaining at each time point was calculated by integrating the area under the curve of the peptide peak at the initial time (t = 0) as a reference. The experiments were done in three replicates.

### Skin abscess infection mouse model

Six-week-old female CD-1 mice were anesthetized, and their backs were shaved. The area was then wiped with ethanol to eliminate any endogenous S*taphylococci* prior to creating a surface abrasion using a scalpel. An aliquot of *S. aureus* ATCC 12600 (10⁷ CFU mL^-1^; 20 μL), grown to an optical density of 0.45 at 600 nm in LB medium and washed twice with sterile PBS (pH 7.4) by centrifugation (5,000 g for 5 minutes), was applied to the scratched region. Peptide or gentamicin, each diluted to the 10×MIC100, was administered to the wound site 2 hours post-infection. One day after the infection, the animals were euthanized, and the infected skin was excised. The tissues were weighed and processed into a homogenized mixture using a bead beater at 25 Hz for 20 minutes. The samples were then serially diluted 10-fold and plated on Mannitol Salt Agar (MSA) plates for colony-forming unit (CFU) quantification. Ten mice per group were included in the study, with two animals designated for histological analysis. All procedures were approved by the University Laboratory Animal Resources (ULAR) at the University of Pennsylvania (Protocol 806763).

### Peritonitis/sepsis mouse model

Six-week-old female CD-1 mice (stock number: 18679700-022) were infected intraperitoneally (I.P.) with *S. aureus* ATCC 12600 (5 × 10^10^ CFU mL^-1^; 100 μL). The bacteria were cultured to an optical density of 0.5 at 600 nm in LB medium, washed twice with sterile PBS (pH 7.4) by centrifugation (3,500 g for 10 minutes), and resuspended in PBS. Mice were treated with three intraperitoneal doses of either gentamicin (460 μmol L^-1^) or peptide (975 μmol L^-1^), each administered in a 100 μL volume at 1.5-hour intervals, beginning 2 hours post-infection. Twenty-four hours after the final treatment, the mice were euthanized, and peritoneal fluids were collected by washing the peritoneal cavity with 5 mL of ice-cold PBS. The bacterial load in the fluids was assessed by dividing the sample into two equal fractions. For extracellular bacteria quantification, one fraction was centrifuged at 300 g for 10 minutes at 4 °C, and the supernatant was plated on MSA plates. For intracellular bacteria quantification, the second fraction was centrifuged at 1,000 rpm for 5 minutes at 4 °C, and the resulting pellet was treated with gentamicin (500 μg mL^-1^) for 3 hours at 37 °C to eliminate extracellular bacteria. The pellet was then washed twice with PBS, serially diluted, and plated on MSA plates. Liver, kidneys, spleen, lungs, and heart were harvested, weighed, and homogenized using a bead beater (25 Hz for 20 minutes). The homogenates were plated on MSA to evaluate bacterial dissemination to these organs. Nine mice per group were included in the study. All procedures were approved by the University Laboratory Animal Resources (ULAR) at the University of Pennsylvania (Protocol 807530).

### Histological analysis

Skin tissues from mice, both untreated and treated with Cy5-Turingcin-46, were collected, fixed in 4% paraformaldehyde in PBS overnight at 4 °C, and stored in 70% ethanol prior to sectioning. Tissue sections were deparaffinized and rehydrated through a series of 5-minute incubations in xylene followed by a graded ethanol series (100% to 70%).

The slides were washed in PBS, and antigen retrieval was performed in citrate buffer (10 mmol L^-1^ citrate, pH 6) at 90 °C for 1 hour. After antigen retrieval, the slides were washed again in PBS and blocked with 2% BSA and 0.05% Tween 20 in PBS for 30 minutes. Sections were then incubated with WGA 488 (10 µg mL^-1^) in 2% BSA and 0.05% Tween 20 in PBS for 30 minutes. Following this, the sections were washed twice in PBS for 3 minutes each and stained with Hoechst 33342 (1 µg mL^-1^) in PBS for 5 minutes. The sections were then washed twice in PBS for 3 minutes each and mounted using a mounting medium. Images were acquired using Zeiss LSM 980 confocal maintained by the CDB Microscopy Core.

### Quantification and statistical analysis Reproducibility of the experimental assays

Unless specified otherwise, all assays were performed in three independent biological replicates, as outlined in the figure legends and the Experimental Models and Methods sections. Cytotoxic and hemolytic activity values were calculated using non-linear regression analysis of peptide screenings across a concentration gradient, representing the concentrations required to kill 50% of the cells in the experiment. For the skin abscess and peritonitis/sepsis infection mouse models, 10 mice per group, including 2 used for histological analysis, and 9 mice per group, respectively, were employed, in accordance with protocols approved by the University Laboratory of Animal Resources (ULAR) at the University of Pennsylvania.

### Statistical tests

In the intracellular antimicrobial activity assays and mouse experiments, all raw data were transformed using a log10 scale. Statistical significance at each time point was assessed independently using one-way ANOVA, followed by Dunnett’s test for intracellular antimicrobial activity and Tukey’s test for mouse experiments. P-values are reported for each group, with all groups compared to the untreated control group.

### Statistical analysis

All calculation and statistical analyses of the experimental data were conducted using GraphPad Prism v.10.3. Statistical significance between different groups was calculated using the tests indicated in each legend.

## Supplementary Information

Figures S1 to S11

Tables S1 to S4

References

