## Supplementary Information for "Design of multimodal antibiotics using deep learning"

**Supporting Information file contains:**

Figures S1 to S11
Tables S1 to S4
References

### Supplementary Figures

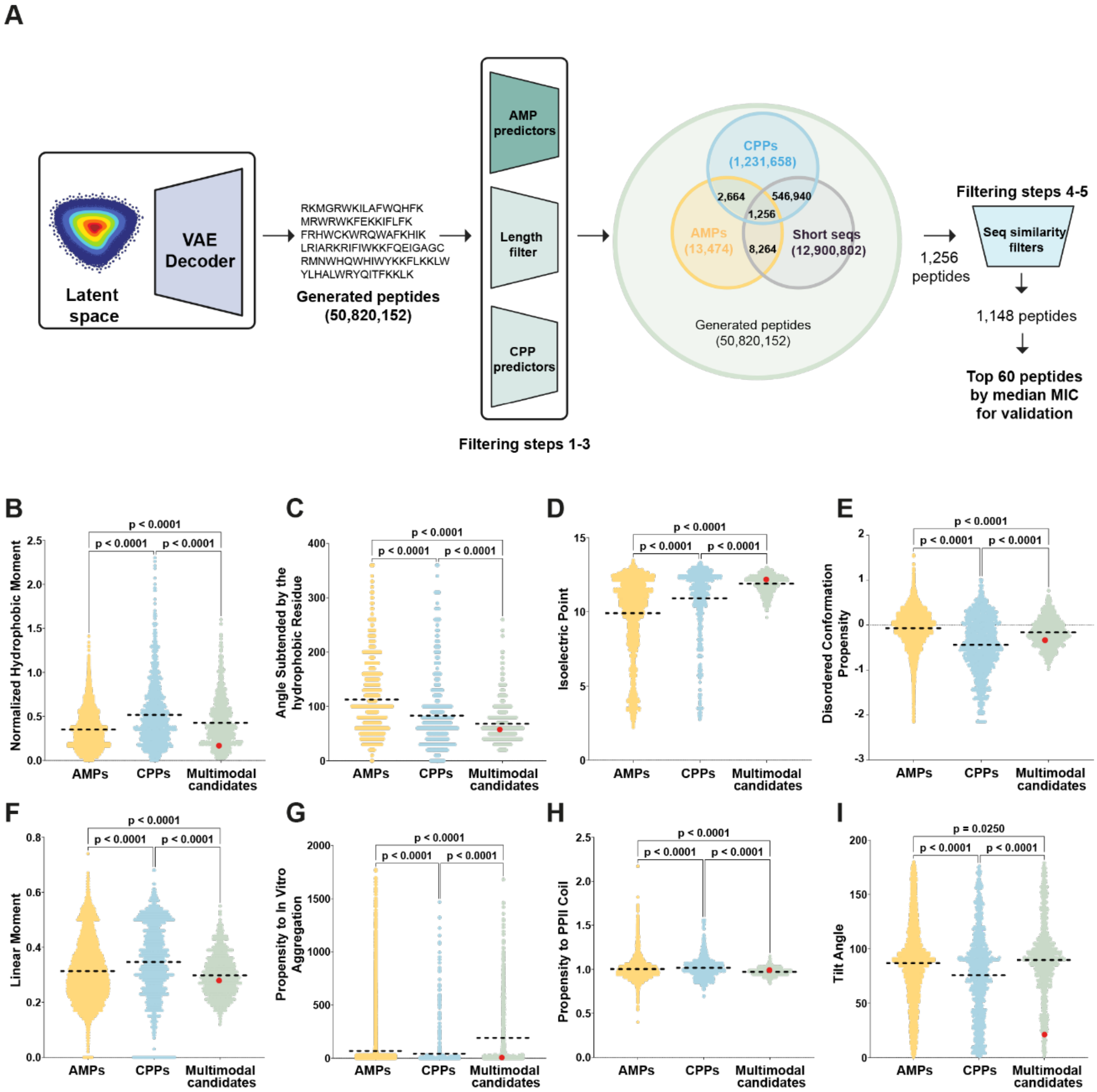


**Figure S1. Deep learning pipeline for multimodal peptide discovery and physicochemical characterization of Turingcins. (A)** Schematic overview of the deep learning framework used to generate peptides with both antimicrobial and cell-penetrating properties. A variational autoencoder (VAE) was used to design novel peptide sequences, followed by filtering through predictive models and similarity checkers to select candidates with high intracellular antimicrobial potential. **(B–I)** Comparative analysis of physicochemical properties of Turingcins versus publicly available antimicrobial peptides (AMPs) and cell-penetrating peptides (CPPs). **(B)** Normalized hydrophobic moment per residue, reflecting how peptides interact with bacterial membranes. **(C)** Angle subtended by hydrophobic residues, influencing membrane binding and insertion. **(D)** Isoelectric point, critical for electrostatic interactions with biological targets. **(E)** Propensity for disordered conformation, associated with membrane perturbation mechanisms. **(F)** Linear moment, influencing peptide binding affinity to biological molecules. **(G)** Propensity for *in vitro* aggregation, affecting peptide self-assembly and potential toxicity. **(H)** Tendency to form a PPII coil, a secondary structure element affecting peptide stability and membrane interactions. **(I)** Tilt angle, determining peptide insertion depth and orientation within the lipid bilayer. All physicochemical properties of the peptides were obtained using the DBAASP server[2]. Statistical significance was assessed using two-tailed t-tests, followed by the Mann-Whitney test; p-values are indicated in the graph, and the dashed line within each distribution represents the mean value for each group. The most promising multimodal candidate from this study, Turingcin-46, is highlighted with a red dot.

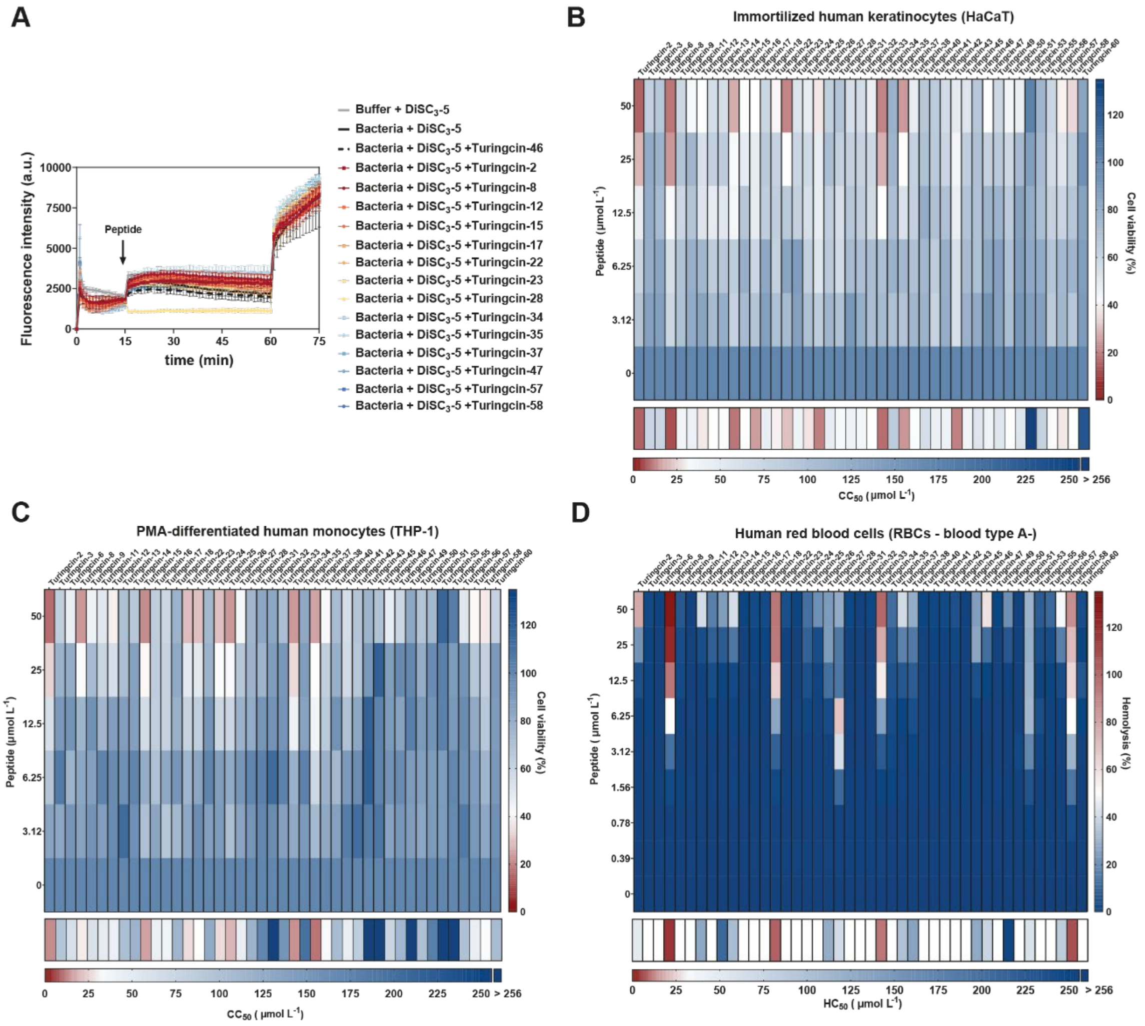


**Figure S2. Mechanism of action and cytotoxicity of Turingcins. (A)** Membrane depolarization effects of Turingcins active against *S. aureus* ATCC 12600, assessed via DiSC3-5 fluorescence assays. Data are presented as raw fluorescence values, with error bars indicating standard deviation from three independent replicates. **(B–D)** Heat maps summarizing cytotoxicity profiles of Turingcins on human immortalized keratinocytes (HaCaT), PMA-differentiated human monocytes (THP-1), and human red blood cells (RBCs). Values represent mean cell viability (%) or hemolysis (%) from three independent experiments. Below each graph, a summary is provided for the predicted $CC_{50}$ or $HC_{50}$ concentrations (µmol $L^{-1}$) of each peptide, indicating the concentration at which 50% of cell death occurs. Values of $CC_{50}$ and $HC_{50}$ were estimated by fitting the dose-response data using a non-linear regression curve.

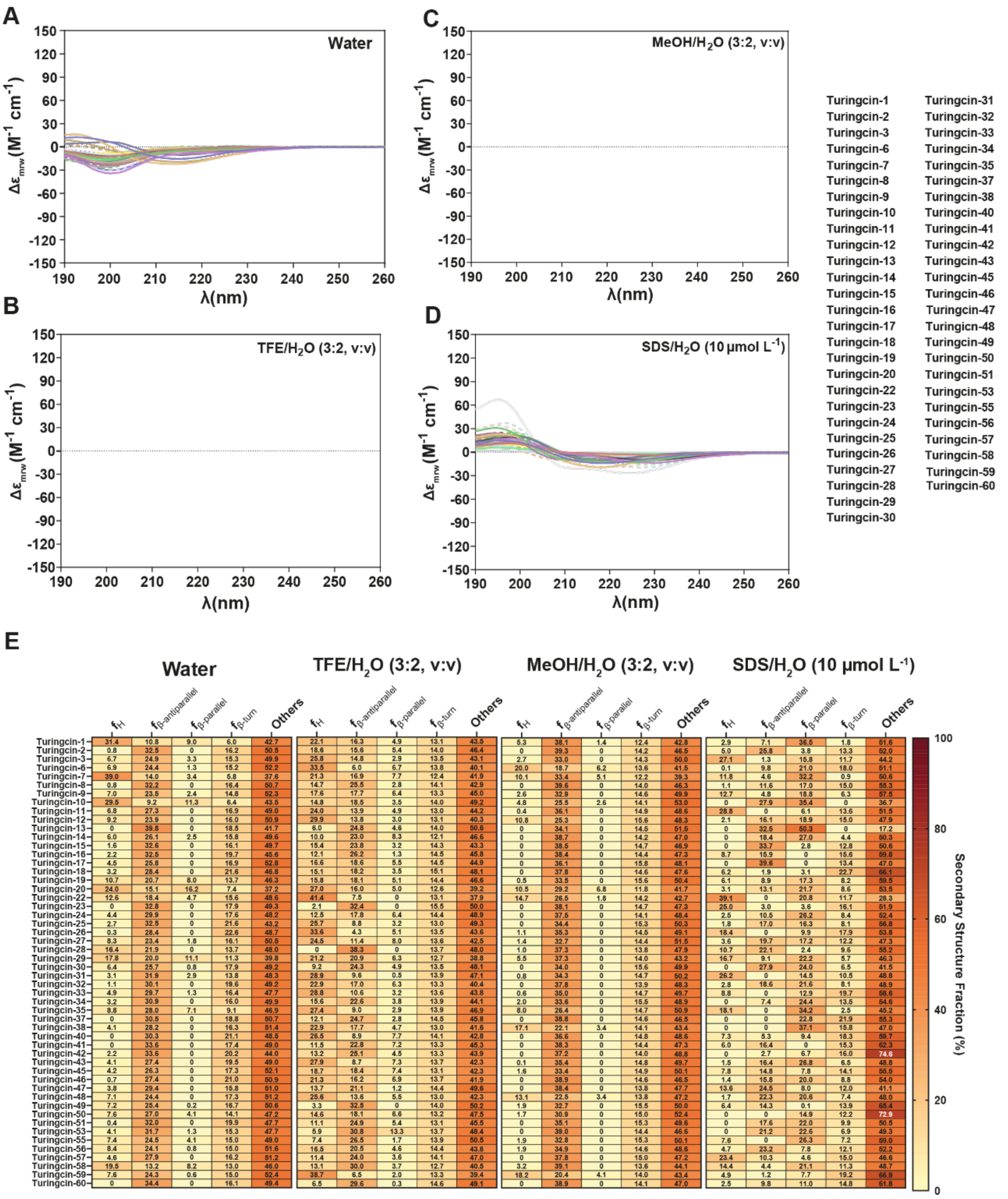


**Figure S3. Secondary structure of Turingcins.** Circular dichroism (CD) spectra for Turingcins were obtained using a J-1500 Jasco CD spectrophotometer. Spectra were recorded in **(A)** water, **(B)** 60% trifluoroethanol in water, **(C)** 60% methanol in water, and **(D)** 10 mmol $L^{-1}$ sodium dodecyl sulfate (SDS) in water. Experiments were conducted at 25 °C in a 1 mm path length quartz cell, with three accumulations captured from 260 to 190 nm at a scan speed of 50 nm $min^{-1}$ and a bandwidth of 0.5 nm. Peptide concentration for all samples was maintained at 50 µmol $L^{-1}$. Heatmap showing the percentage of secondary structure present in each peptide across three different solvents: water, 60% trifluoroethanol in water, 60% methanol in water, and SDS in water (10 mmol $L^{-1}$). The secondary structure fractions were calculated using the BeStSel server [1].

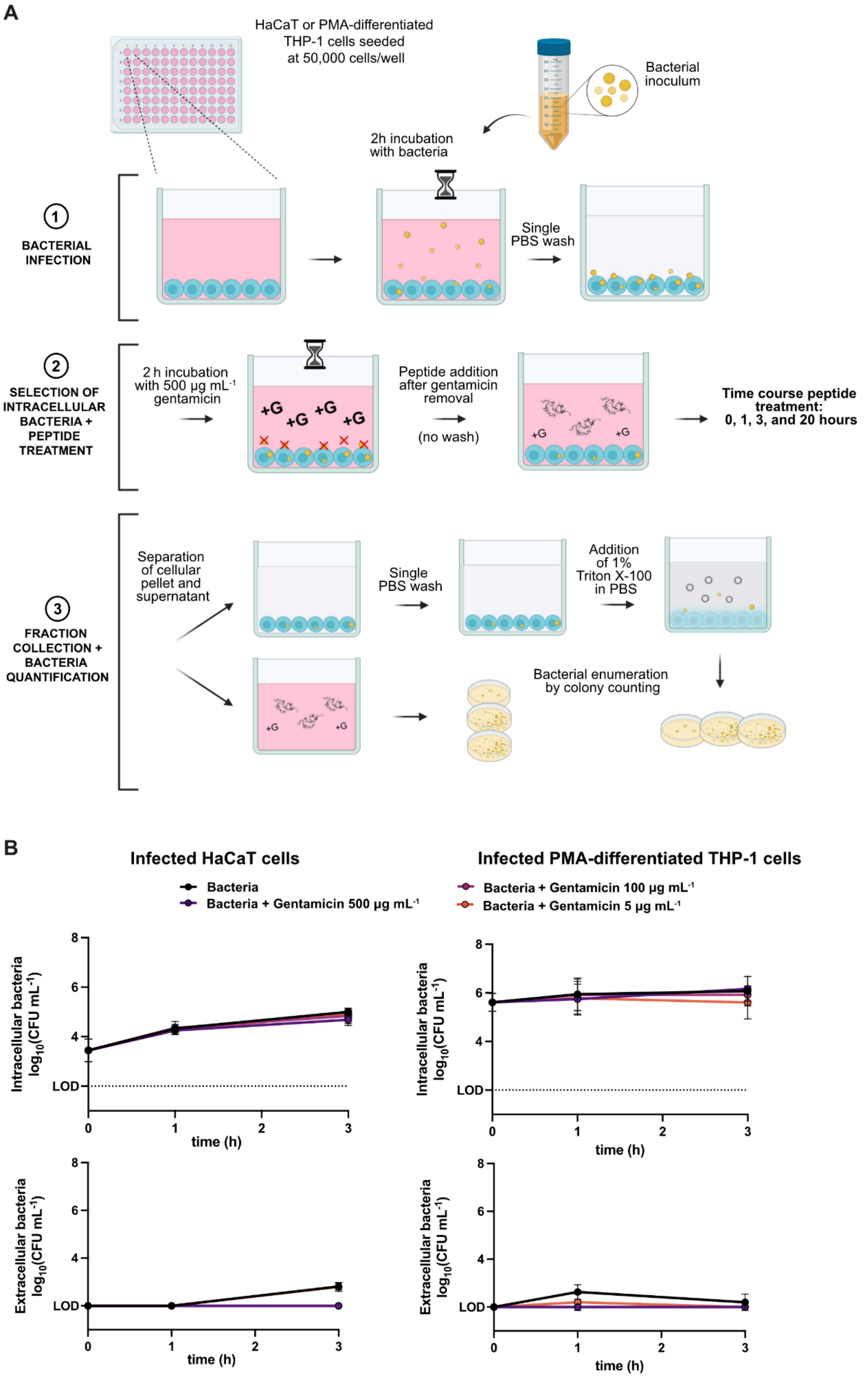
A
HaCaT or PMA-differentiated THP-1 cells seeded at 50,000 cells/well
Bacterial inoculum
2h incubation with bacteria
1 BACTERIAL INFECTION
Single PBS wash
2 SELECTION OF INTRACELLULAR BACTERIA + PEPTIDE TREATMENT
2 h incubation with 500 μg mL-1 gentamicin
+G
Peptide addition after gentamicin removal
(no wash)
Time course peptide treatment: 0, 1, 3, and 20 hours
3 FRACTION COLLECTION + BACTERIA QUANTIFICATION
Separation of cellular pellet and supernatant
Single PBS wash
Addition of 1% Triton X-100 in PBS
Bacterial enumeration by colony counting
B
Infected HaCaT cells
Infected PMA-differentiated THP-1 cells
Bacteria
Bacteria + Gentamicin 500 μg mL-1
Bacteria + Gentamicin 100 μg mL-1
Bacteria + Gentamicin 5 μg mL-1
Intracellular bacteria log10(CFU mL-1)
Extracellular bacteria log10(CFU mL-1)
LOD
time (h)

**Figure S4. Intracellular antimicrobial activity. (A)** Schematic overview of the experimental procedure for assessing intracellular antimicrobial activity, illustrating three main steps: bacterial infection, selection of intracellular bacteria followed by peptide treatment, and fraction collection with subsequent bacterial quantification. **(B)** Time course of gentamicin activity against intracellular *S. aureus* ATCC 12600 in HaCaT keratinocytes and PMA-differentiated THP-1 macrophages. Intracellular bacterial loads were measured at 0, 1, and 3 hours after treatment with gentamicin at 5, 100, and 500 µg $mL^{-1}$. Data represent three independent experiments for each condition.

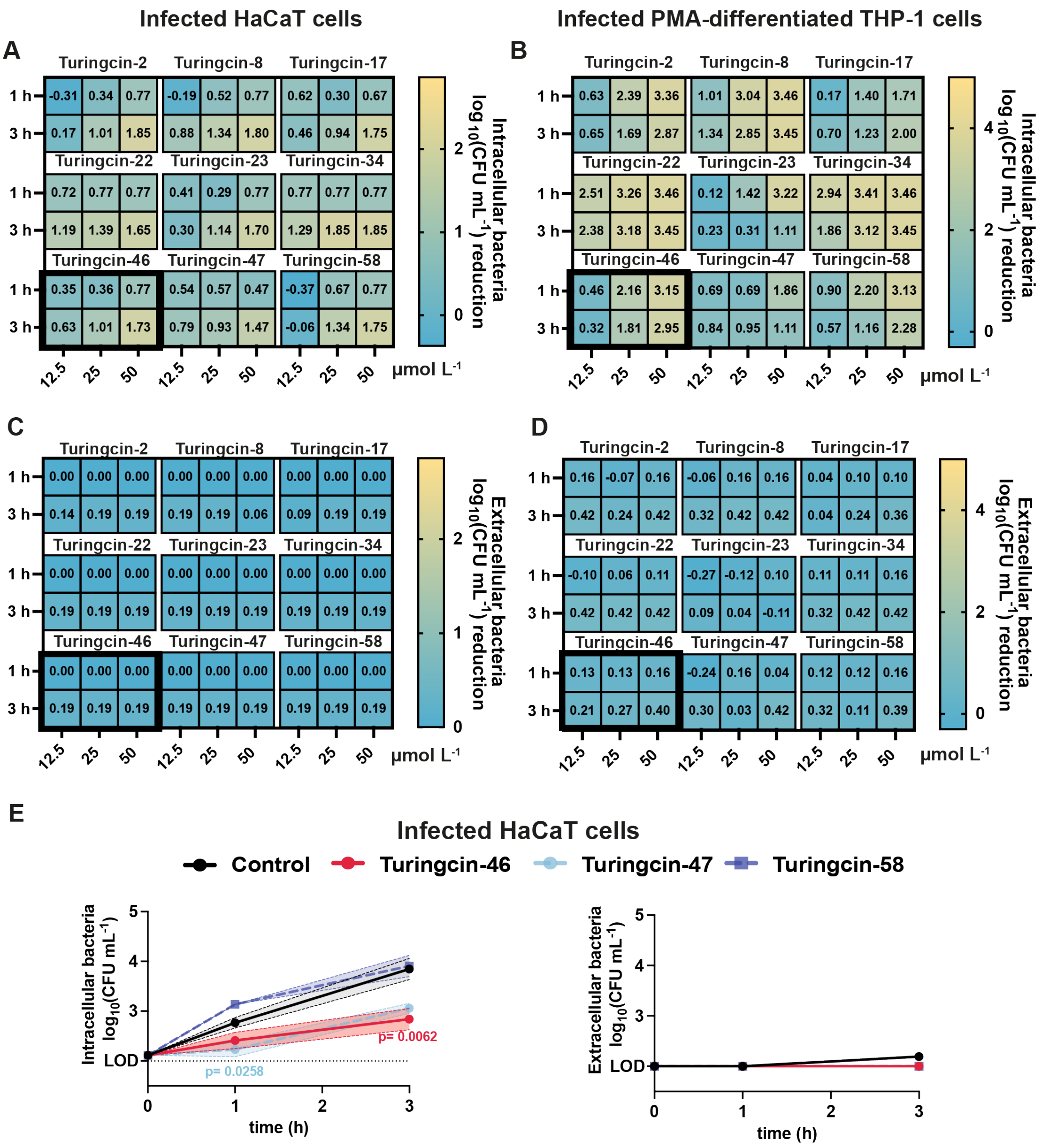


**Figure S5. Intracellular antimicrobial activity of Turingcins against *S. aureus*.** Time-course evaluation of intracellular **(A, B)** and extracellular **(C, D)** antimicrobial activity of Turingcins in *S. aureus* ATCC 12600-infected HaCaT keratinocytes and PMA-differentiated THP-1 macrophages. Bacterial loads were measured at 0, 1, and 3 hours following treatment with Turingcins at 12.5, 25, and 50 μmol mL$^{-1}$, including concentrations known to elicit cytotoxic effects. The $\log_{10}$(CFU mL$^{-1}$) reduction represents the difference in bacterial load between treated and untreated groups at each time point. A minimum of two independent experiments was performed for all conditions, with the lead compound (Turingcin-46) tested in three. **(E)** Time-course analysis (0, 1, and 3 hours) of intracellular and extracellular antimicrobial activity of Turingcin peptides at non-cytotoxic concentrations against *S. aureus* ATCC 12600 in HaCaT keratinocytes.

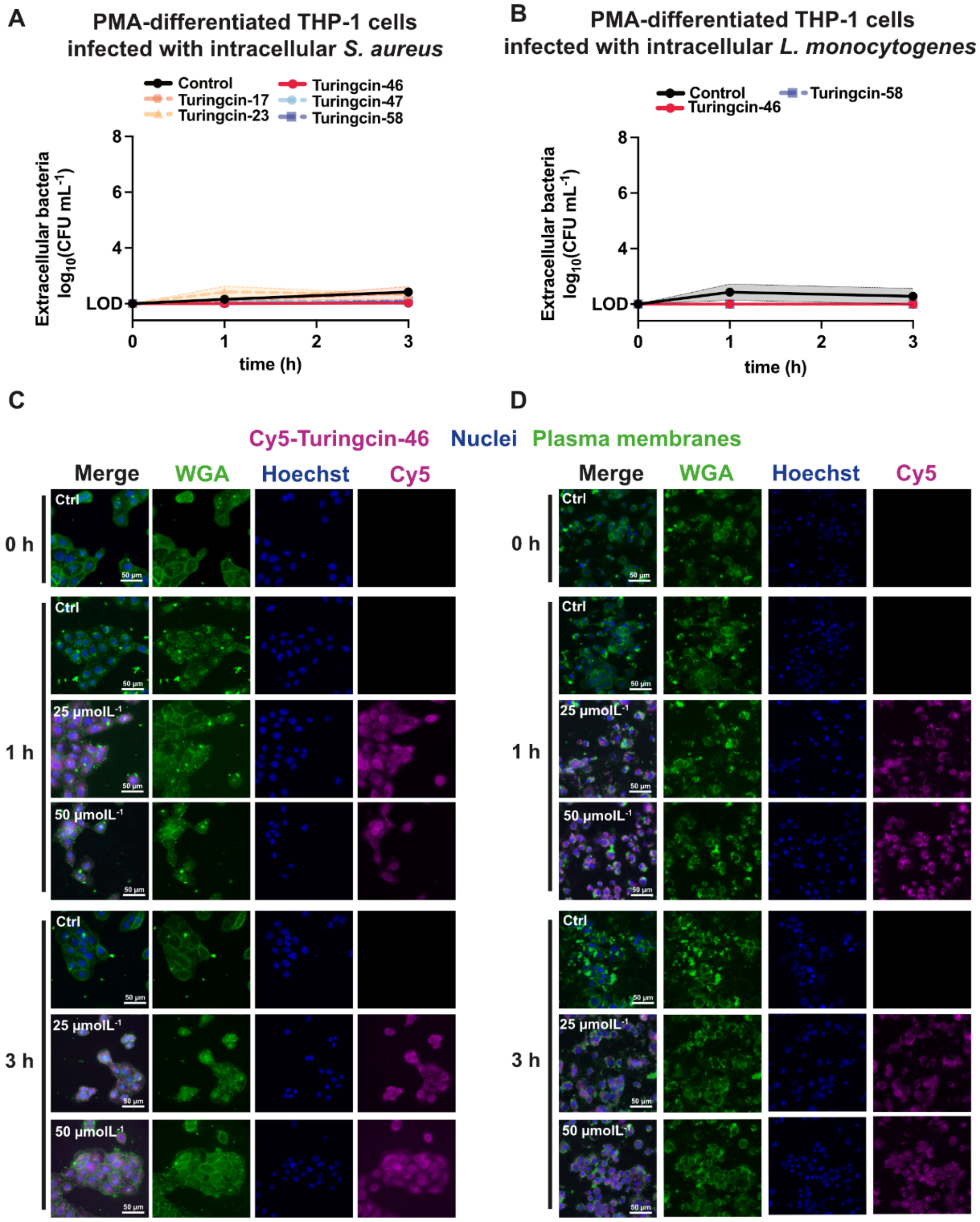


**Figure S6. Intracellular antimicrobial activity of Turingcins.** Time-course evaluation of the extracellular antimicrobial activity of Turingcin peptides in **(A)** *S. aureus* ATCC 1600-infected and **(B)** *Listeria monocytogenes* ATCC 19111-infected PMA-differentiated THP-1 macrophages. Bacterial loads were measured at 0, 1, and 3 hours following treatment with Turingcins at non-cytotoxic concentrations. **(C, D)** Representative fluorescence microscopy images of HaCaT and PMA-differentiated THP-1 cells infected with *S. aureus* ATCC 12600 and treated with Cy5-Turingcin-46 at 0, 1, and 3-hours post-treatment. Cells were fixed with 4% paraformaldehyde and stained with WGA 488 (10 µg mL$^{-1}$) to visualize plasma membranes and Hoechst 33342 (1 µg mL$^{-1}$) for nuclei. Brighter green puncta in the WGA channel correspond to bacterial aggregates. These appear more intense due to clustering of WGA-positive bacteria, which enhances the fluorescence signal. Each experiment was performed in biological duplicate, with two technical replicates each.

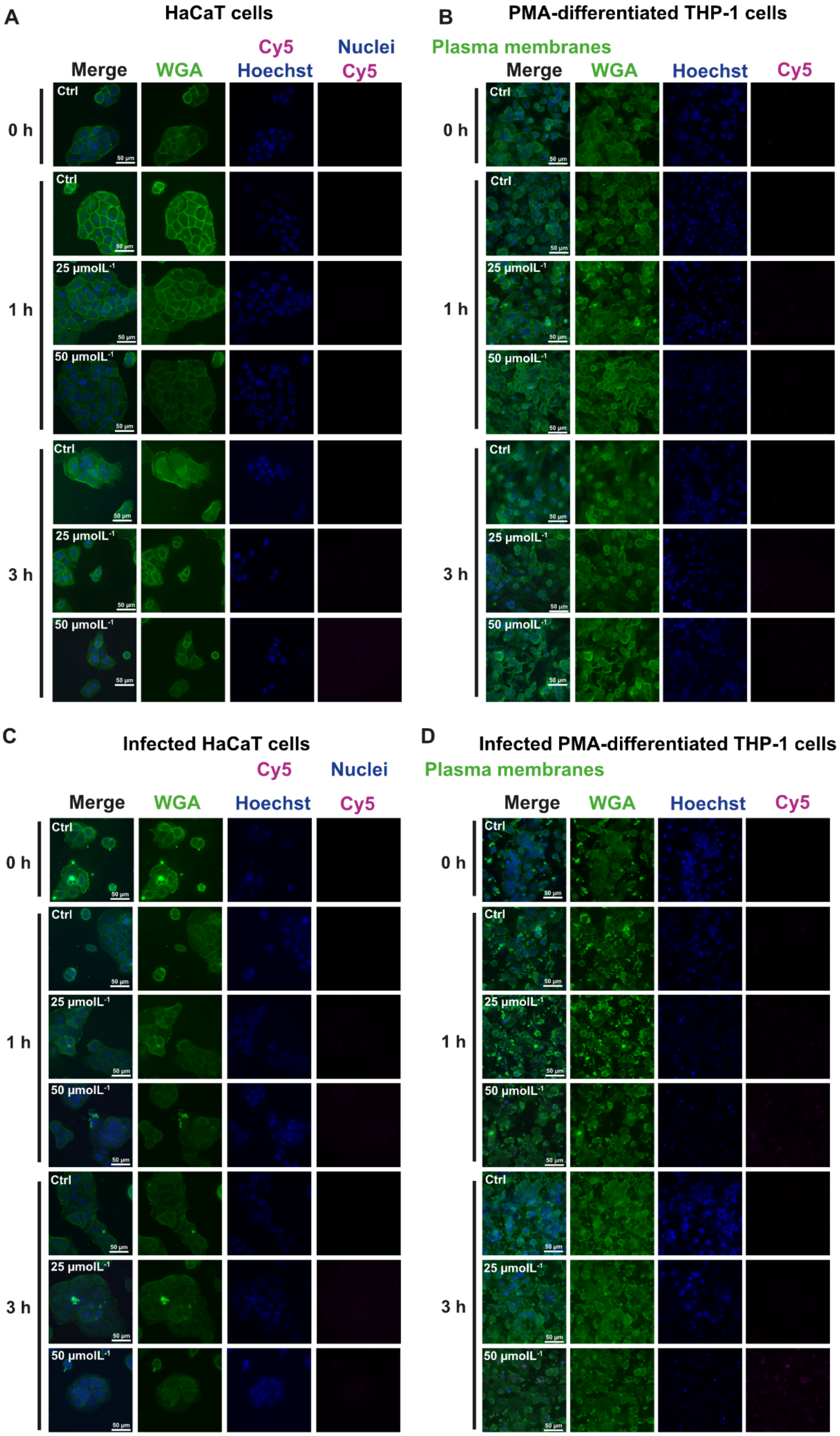
A
HaCaT cells
Cy5 Hoechst
Nuclei Cy5
Merge
WGA
0 h
1 h
3 h
Ctrl
25 μmolL-1
50 μmolL-1
50 μm
B
PMA-differentiated THP-1 cells
Plasma membranes
Merge
WGA
Hoechst
Cy5
C
Infected HaCaT cells
D
Infected PMA-differentiated THP-1 cells

**Figure S7. Control uptake and localization of Cy5 alone.** Representative fluorescence microscopy images of **(A, B)** uninfected and **(C, D)** *S. aureus*-infected HaCaT keratinocytes and PMA-differentiated THP-1 macrophages treated with Cy5 at 0, 1, and 3 hours post-treatment. Cells were fixed with 4% paraformaldehyde and stained with WGA-Alexa Fluor 488 (10 $\mu g\ mL^{-1}$) to visualize plasma membranes, and Hoechst 33342 (1 $\mu g\ mL^{-1}$) to stain nuclei. Each experiment was performed in biological duplicate with two technical replicates. Brighter green puncta in the WGA channel correspond to bacterial aggregates. These appear more intense due to clustering of WGA-positive bacteria, which enhances the fluorescence signal.

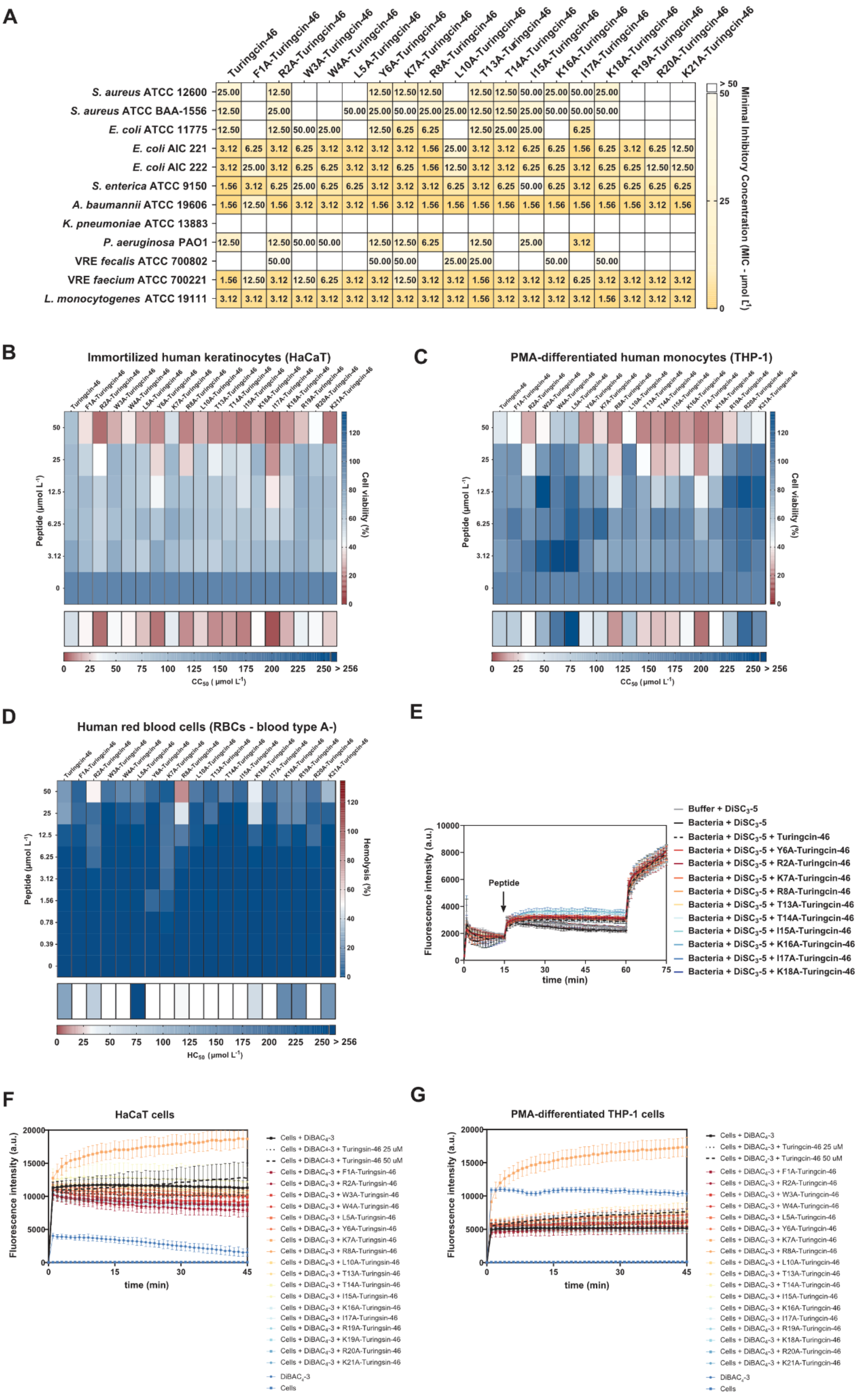
A
S. aureus ATCC 12600
S. aureus ATCC BAA-1556
E. coli ATCC 11775
E. coli AIC 221
E. coli AIC 222
S. enterica ATCC 9150
A. baumannii ATCC 19606
K. pneumoniae ATCC 13883
P. aeruginosa PAO1
VRE fecalis ATCC 700802
VRE faecium ATCC 700221
L. monocytogenes ATCC 19111
Minimal Inhibitory Concentration (MIC - μmol L-1)
B
Immortilized human keratinocytes (HaCaT)
Peptide (μmol L-1)
Cell viability (%)
CC50 (μmol L-1)
C
PMA-differentiated human monocytes (THP-1)
D
Human red blood cells (RBCs - blood type A-)
Hemolysis (%)
HC50 (μmol L-1)
E
Fluorescence intensity (a.u.)
time (min)
Peptide
Buffer + DiSC3-5
Bacteria + DiSC3-5
Bacteria + DiSC3-5 + Turingcin-46
F
HaCaT cells
G
PMA-differentiated THP-1 cells
Cells + DiBAC4-3
DiBAC4-3
Cells

**Figure S8. Antimicrobial activity, cytotoxicity and mechanism of action of Turingcin-46 and its alanine-substituted derivatives. (A)** Heat map showing the antimicrobial activity (MIC values in µmol $L^{-1}$) of alanine-substituted Turingcin-46 variants against clinically relevant bacterial strains, including antibiotic-resistant species. Bacterial cultures were exposed to peptide concentrations ranging from 0 to 50 µmol $L^{-1}$ at 37°C, and growth inhibition was assessed after 24 hours by measuring optical density at 600 nm. MIC values represent the most frequently observed results across three experimental replicates**. (B–D)** Heat maps illustrating the cytotoxic effects of Turingcin-46 and its Ala-derivatives on human immortalized keratinocytes (HaCaT), PMA-differentiated human monocytes (THP-1), and human red blood cells (RBCs). The color scales represent cell viability (%) or hemolysis (%), derived from the mean values obtained in three experiments. A summary of the predicted $CC_{50}$ or $HC_{50}$ concentrations (µmol $L^{-1}$) for each peptide is shown beneath each graph, representing the concentration required to induce 50% cell death. These values were determined by applying a non-linear regression curve to the dose-response data. **(E)** Membrane depolarization of *S. aureus* ATCC 12600 upon treatment with Turingcin-46 and eight active Ala-derivatives was assessed using the voltage-sensitive dye $DiSC_3$-5. Raw fluorescence intensity values are presented with error bars indicating standard deviations from three independent replicates. **(F-G)** The impact on eukaryotic cell membrane potential, measured as depolarization or hyperpolarization, was assessed for HaCaT and PMA-differentiated THP-1 cells treated with Turingcin-46 and its alanine variants. The evaluation was performed using the $DiBAC_4$-3 dye at $CC_{50}$ concentrations. Raw fluorescence intensity values are shown, with error bars representing the standard deviation from three independent replicates.

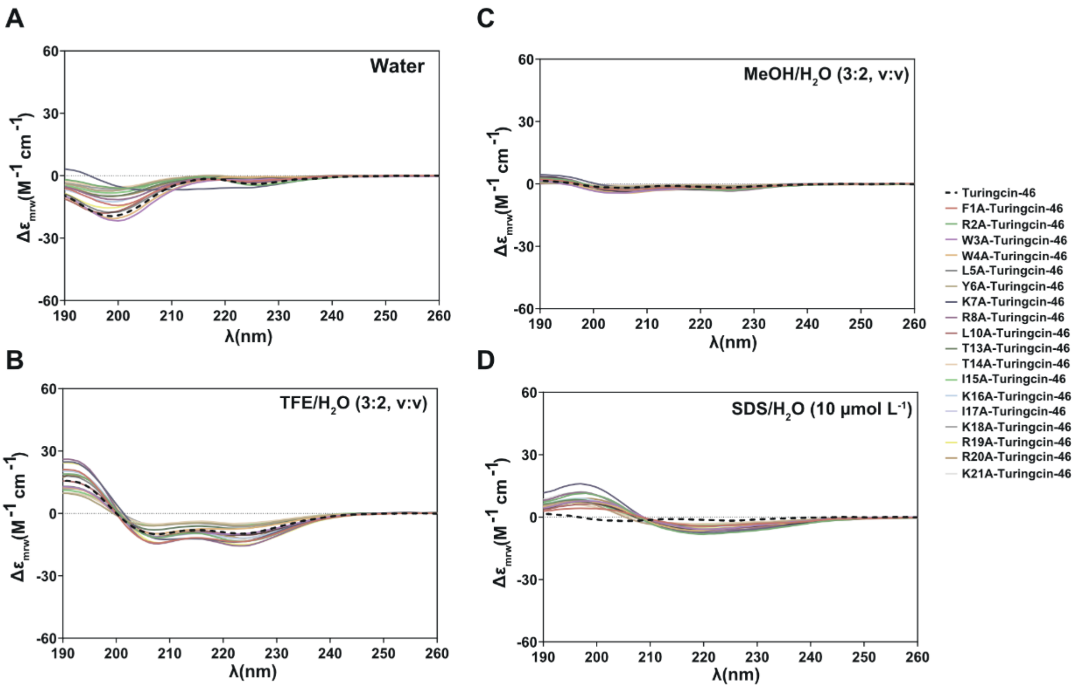


E

Secondary Structure Fraction (%)

| | Water $f_H$ | Water $f_{\beta\text{-antiparallel}}$ | Water $f_{\beta\text{-parallel}}$ | Water $f_{\beta\text{-turn}}$ | Water Others | TFE/$H_2O$ (3:2, v:v) $f_H$ | TFE/$H_2O$ $f_{\beta\text{-antiparallel}}$ | TFE/$H_2O$ $f_{\beta\text{-parallel}}$ | TFE/$H_2O$ $f_{\beta\text{-turn}}$ | TFE/$H_2O$ Others |
|---|---|---|---|---|---|---|---|---|---|---|
| Turingcin-46 | 0.7 | 27.4 | 0 | 21.0 | 50.9 | 21.3 | 16.2 | 6.9 | 13.7 | 41.9 |
| F1A-Turingcin-46 | 0 | 29.6 | 0 | 17.7 | 52.8 | 31.5 | 6.9 | 6.6 | 13.8 | 41.2 |
| R2A-Turingcin-46 | 0 | 29.3 | 0 | 18.5 | 52.2 | 30.5 | 3.0 | 7.3 | 16.1 | 43.1 |
| W3A-Turingcin-46 | 3.1 | 25.4 | 0 | 19.6 | 52.0 | 25.8 | 13.0 | 7.8 | 13.7 | 39.7 |
| W4A-Turingcin-46 | 2.0 | 29.2 | 0 | 20.6 | 48.3 | 19.6 | 19.7 | 5.7 | 13.6 | 41.4 |
| L5A-Turingcin-46 | 0 | 31.8 | 0 | 18.7 | 49.5 | 28.0 | 15.2 | 3.6 | 14.0 | 39.1 |
| Y6A-Turingcin-46 | 0 | 36.0 | 0 | 16.9 | 47.1 | 11.3 | 26.3 | 1.6 | 14.0 | 46.8 |
| K7A-Turingcin-46 | 12.2 | 24.6 | 5.4 | 13.9 | 44.0 | 29.9 | 15.2 | 6.1 | 11.8 | 37.1 |
| R8A-Turingcin-46 | 0 | 29.0 | 0 | 18.5 | 52.5 | 35.3 | 7.1 | 1.2 | 15.8 | 40.6 |
| L10A-Turingcin-46 | 0 | 30.1 | 0 | 20.1 | 49.8 | 25.0 | 13.1 | 5.9 | 14.6 | 41.5 |
| T13A-Turingcin-46 | 0 | 32.2 | 0 | 16.9 | 50.9 | 15.6 | 23.5 | 4.3 | 13.4 | 43.2 |
| T14A-Turingcin-46 | 0 | 35.5 | 0 | 16.0 | 48.6 | 8.5 | 29.5 | 0 | 14.4 | 47.5 |
| I15A-Turingcin-46 | 0 | 33.8 | 0 | 16.9 | 49.3 | 9.5 | 29.2 | 0 | 15.1 | 46.2 |
| K16A-Turingcin-46 | 1.3 | 28.1 | 0 | 18.1 | 52.5 | 28.1 | 11.8 | 5.8 | 14.2 | 40.1 |
| I17A-Turingcin-46 | 0 | 33.8 | 0 | 16.0 | 50.2 | 9.9 | 28.7 | 0.3 | 15.3 | 45.8 |
| K18A-Turingcin-46 | 0 | 34.9 | 0 | 16.4 | 48.7 | 11.5 | 27.8 | 1.5 | 14.0 | 45.2 |
| R19A-Turingcin-46 | 0.4 | 27.7 | 0 | 20.0 | 52.0 | 37.1 | 2.5 | 5.6 | 15.4 | 39.5 |
| R20A-Turingcin-46 | 0 | 30.8 | 0 | 17.7 | 51.6 | 26.1 | 13.3 | 3.0 | 15.7 | 42.0 |
| K21A-Turingcin-46 | 0 | 29.0 | 0 | 18.4 | 52.6 | 29.4 | 7.6 | 5.2 | 15.6 | 42.2 |

| | MeOH/$H_2O$ (3:2, v:v) $f_H$ | MeOH/$H_2O$ $f_{\beta\text{-antiparallel}}$ | MeOH/$H_2O$ $f_{\beta\text{-parallel}}$ | MeOH/$H_2O$ $f_{\beta\text{-turn}}$ | MeOH/$H_2O$ Others | SDS/$H_2O$ (10 μmol $L^{-1}$) $f_H$ | SDS/$H_2O$ $f_{\beta\text{-antiparallel}}$ | SDS/$H_2O$ $f_{\beta\text{-parallel}}$ | SDS/$H_2O$ $f_{\beta\text{-turn}}$ | SDS/$H_2O$ Others |
|---|---|---|---|---|---|---|---|---|---|---|
| Turingcin-46 | 0 | 38.9 | 0 | 14.6 | 46.5 | 1.4 | 15.8 | 20.0 | 8.8 | 54.0 |
| F1A-Turingcin-46 | 0 | 37.9 | 0 | 15.2 | 46.9 | 2.4 | 28.0 | 9.9 | 11.3 | 48.3 |
| R2A-Turingcin-46 | 1.1 | 36.0 | 0 | 14.7 | 48.2 | 2.7 | 19.8 | 22.3 | 9.2 | 46.0 |
| W3A-Turingcin-46 | 0 | 37.1 | 0 | 15.4 | 47.5 | 3.5 | 26.0 | 14.4 | 9.6 | 46.6 |
| W4A-Turingcin-46 | 0.8 | 36.2 | 0 | 15.8 | 47.2 | 2.5 | 24.8 | 16.0 | 9.2 | 47.5 |
| L5A-Turingcin-46 | 1.1 | 37.0 | 0 | 15.1 | 46.9 | 3.5 | 23.5 | 14.4 | 10.1 | 48.5 |
| Y6A-Turingcin-46 | 0 | 39.3 | 0 | 14.4 | 46.3 | 4.4 | 30.6 | 10.2 | 11.0 | 43.8 |
| K7A-Turingcin-46 | 0 | 42.1 | 0 | 13.8 | 44.1 | 1.8 | 22.8 | 20.9 | 7.9 | 46.7 |
| R8A-Turingcin-46 | 0.5 | 37.6 | 0 | 15.3 | 46.6 | 4.4 | 22.8 | 13.0 | 10.9 | 49.0 |
| L10A-Turingcin-46 | 0 | 38.2 | 0 | 14.9 | 46.9 | 1.1 | 24.5 | 16.1 | 10.2 | 48.1 |
| T13A-Turingcin-46 | 0 | 38.7 | 0 | 14.3 | 47.0 | 4.1 | 29.4 | 9.0 | 11.2 | 46.3 |
| T14A-Turingcin-46 | 0 | 40.3 | 0 | 14.5 | 45.2 | 1.7 | 33.1 | 5.6 | 12.1 | 47.5 |
| I15A-Turingcin-46 | 0 | 40.3 | 0 | 14.6 | 45.1 | 5.0 | 26.7 | 8.2 | 11.4 | 48.8 |
| K16A-Turingcin-46 | 1.9 | 36.4 | 0 | 15.3 | 46.4 | 3.9 | 22.0 | 15.5 | 10.8 | 47.8 |
| I17A-Turingcin-46 | 0 | 39.6 | 0 | 14.5 | 45.9 | 4.9 | 26.1 | 9.5 | 11.3 | 48.1 |
| K18A-Turingcin-46 | 0 | 40.3 | 0 | 14.5 | 45.2 | 4.9 | 27.4 | 8.5 | 11.5 | 47.8 |
| R19A-Turingcin-46 | 0 | 38.6 | 0 | 14.5 | 46.9 | 2.3 | 21.4 | 16.6 | 10.3 | 49.3 |
| R20A-Turingcin-46 | 0 | 37.6 | 0 | 14.4 | 48.1 | 4.1 | 23.7 | 10.9 | 11.7 | 49.7 |
| K21A-Turingcin-46 | 0 | 37.5 | 0 | 14.7 | 47.7 | 3.3 | 23.0 | 15.9 | 11.0 | 46.8 |

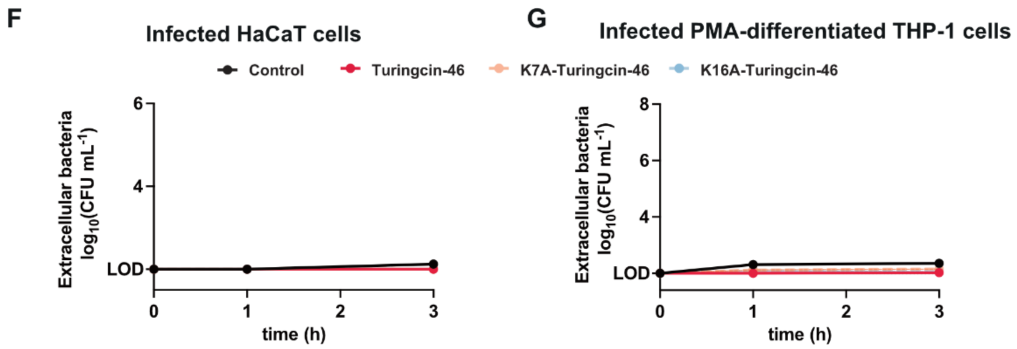

**Figure S9. Secondary structure and intracellular antimicrobial activity of Turingcin-46 and its alanine-substituted variants.** Circular dichroism (CD) spectroscopy was performed to assess the secondary structure of Turingcin-46 and its Ala-derivatives at 50 μmol $L^{-1}$. Measurements were recorded using a J-1500 Jasco CD spectrophotometer at 25 °C in a 1 mm path length quartz cell, with three accumulations collected per sample. Spectra were obtained in four different environments: **(A)** Water; **(b)** 60% trifluoroethanol (TFE) in water; **(C)** 60% methanol (MeOH) in water; **(d)** Sodium dodecyl sulfate (SDS, 10 mmol $L^{-1}$) in water. Scans were conducted from 260 to 190 nm at a speed of 50 nm $min^{-1}$ and a bandwidth of 0.5 nm. **(E)** A heatmap summarizing the secondary structure proportions (α-helix, β-sheet, and disordered) of each peptide in the four tested solvents. Secondary structure fractions were determined using the BeStSel server [1]. **(F, G)** Time-course analysis (0, 1, and 3 hours) of extracellular antimicrobial activity of Turingcin-46, K7A-Turingcin-46, and K16A-Turingcin-46 at non-cytotoxic concentrations in *S. aureus*-infected HaCaT keratinocytes and PMA-differentiated THP-1 macrophages. All experiments were carried out in three independent replicates.

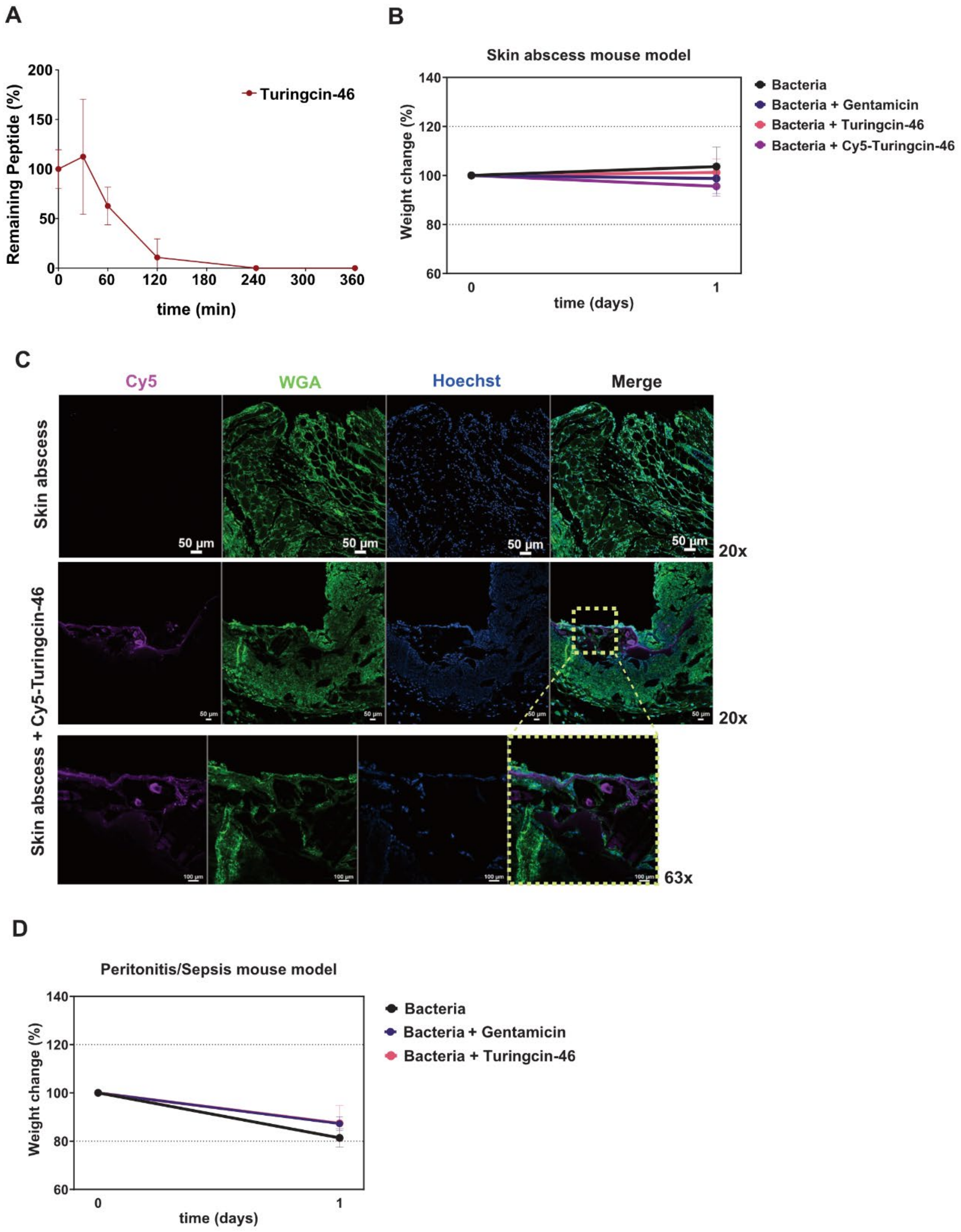


**Figure S10. Stability, body weight monitoring, and histological analysis in murine infection models. (A)** Stability profile of Turingcin-46 exposed to human serum for 6 hours, evaluating peptide resistance to proteolytic degradation. **(B)** Mouse body weight monitoring in the skin abscess model to assess potential toxicity from both the *S. aureus* infection and Turingcin-46 treatment. **(C)** Representative confocal laser scanning microscopy (CLSM) images of excised skin tissues from untreated mice and those treated with Cy5-Turingcin-46. Deparaffinized tissue sections were stained with WGA 488 (10 μg $mL^{-1}$) to highlight plasma membranes and Hoechst 33342 (1 μg $mL^{-1}$) to visualize nuclei. **(D)** Mouse body weight was also monitored in the peritonitis/sepsis infection model to evaluate any potential toxic effects from the bacterial infection and Turingcin-46 treatment.

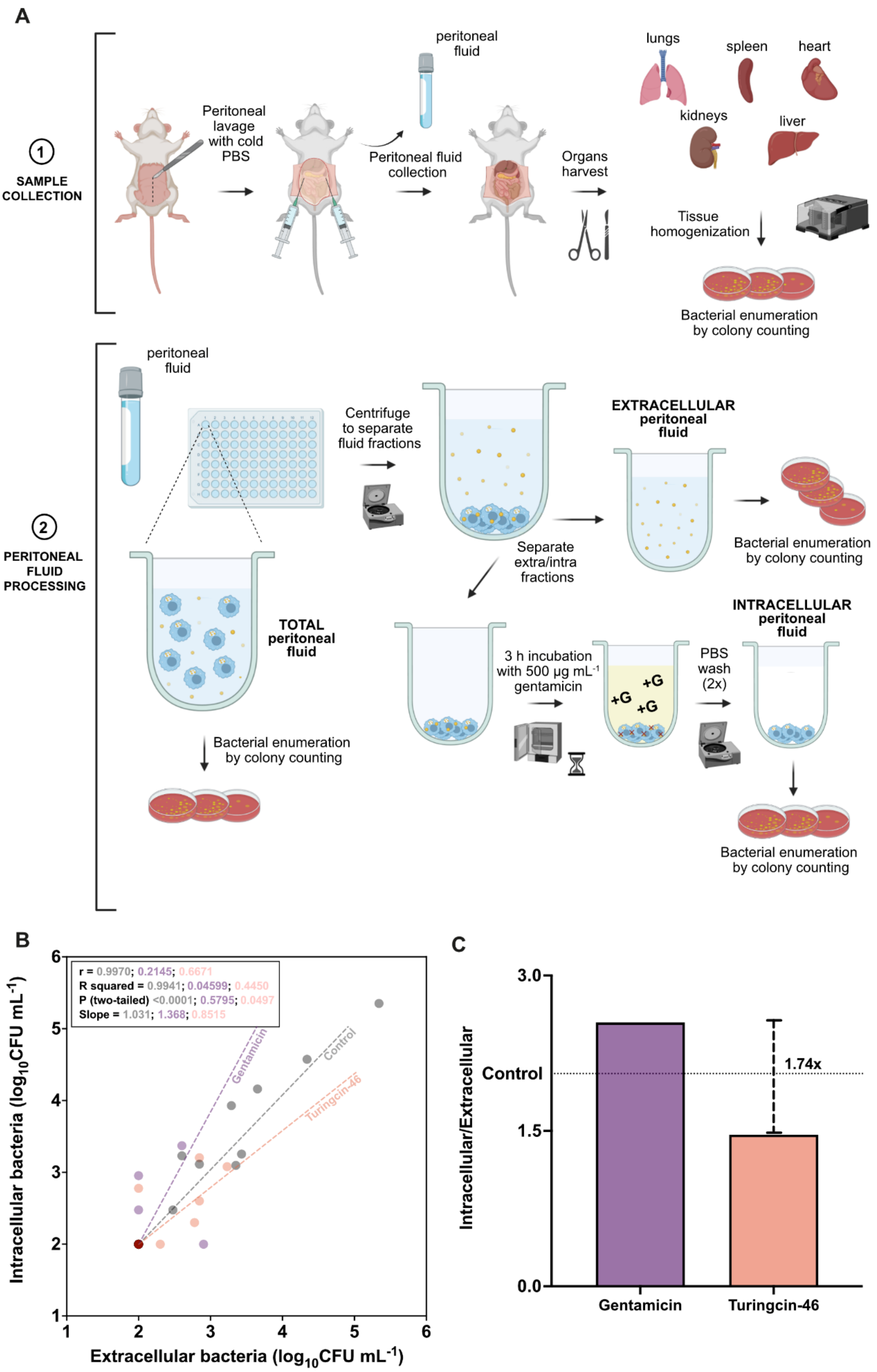


**Figure S11. Anti-infective activity of Turingcin-46 in a murine peritonitis/sepsis model. (A)** Schematic overview of the peritonitis/sepsis mouse model used to collect intracellular and extracellular bacterial loads in peritoneal fluid. **(B)** Correlations between intracellular and extracellular *S. aureus* CFU counts in peritoneal lavage samples from infected mice, plotted on a $\log_{10}$ scale axes. Each point represents an individual mouse (n = 9 per group; total n= 27). Pearson correlation analysis was performed independently for each treatment group. The corresponding correlation coefficient (r), coefficient of determination ($R^2$), two-tailed p-value, and slope are reported. Linear regression lines are shown for each group. **(C)** Bar graph comparing the average ratio of intracellular to extracellular bacterial loads for gentamicin- and Turingcin-46-treated groups over nine replicates.

### Tables

**Table S1.** Selection of 52 soluble Turingcins from 60 top-ranked compounds for experimental testing.

| Peptide | Sequence | Solubility in autoclaved deionized water |
|---|---|---|
| Turingcin-1 | RRWVHKILLIIIKILA | Soluble |
| Turingcin-2 | RLIKAILNRVWKAWAIQKK | Soluble |
| Turingcin-3 | KKVWYIKVKLALALIKNI | Soluble |
| Turingcin-4 | ILARLSSILKLI | Insoluble |
| Turingcin-5 | KILWYRIILIRFQKVW | Insoluble |
| Turingcin-6 | KRRAARIKQLRFIQQSLREERIIQKLIK | Soluble |
| Turingcin-7 | RLKSILLLIVTKINIRLKIA | Soluble |
| Turingcin-8 | WAVKSSRIIYTLLRKLKTWFNIKAK | Soluble |
| Turingcin-9 | RILRKVWTAWLREAALTIKVKR | Soluble |
| Turingcin-10 | RYIRLLVILKLA | Soluble |
| Turingcin-11 | RWWYTSIIKKKLNKIIA | Soluble |
| Turingcin-12 | KILIRKILTIIERKLLRSNLKS | Soluble |
| Turingcin-13 | KMWKLRIKLTI | Soluble |
| Turingcin-14 | KIWKIKILRINKKTSSIIA | Soluble |
| Turingcin-15 | SRKYRQLAKYIWRNLANKNRLIIKA | Soluble |
| Turingcin-16 | KRIKILRKYKWNQQYKKKL | Soluble |
| Turingcin-17 | LIWRRKIFRIKLRI | Soluble |
| Turingcin-18 | KKRVYRQKIRLIKKAIFFKKR | Soluble |
| Turingcin-19 | KKIIIKFLIFNTKL | Soluble |
| Turingcin-20 | RRLNSKINLYLRIIIALINK | Soluble |
| Turingcin-21 | WYLIRVALIAFKKA | Insoluble |
| Turingcin-22 | RIINIILNRNLKKIILAAAIRNAIRW | Soluble |
| Turingcin-23 | VMRRFFRLLRKIKL | Soluble |
| Turingcin-24 | KQIKQRSIRIKIFKKIL | Soluble |
| Turingcin-25 | RVWKTLRIKVYFARRKQR | Soluble |
| Turingcin-26 | KLRSSKKIFQIAAKFNKLFRRK | Soluble |
| Turingcin-27 | QRRVLRFKLNTKIARLINLNI | Soluble |
| Turingcin-28 | RILRKWSNWWWNNWLIWK | Soluble |

| | | |
|---|---|---|
| Turingcin-29 | RINIIIHIAWRSKLIL | Soluble |
| Turingcin-30 | KILFLFITTKKL | Soluble |
| Turingcin-31 | TKLWAIINKALRKI | Soluble |
| Turingcin-32 | RKYKISNRIRIIEKKKRLFRNQ | Soluble |
| Turingcin-33 | RLRGGGKLRLLHKIVLNNRII | Soluble |
| Turingcin-34 | RTAAYGKYLKRWLKKMIVKWK | Soluble |
| Turingcin-35 | RRIVNRISVWTKLALLTIKMI | Soluble |
| Turingcin-36 | RLAVTIITAKLALNSNWNLIRKALWRI | Insoluble |
| Turingcin-37 | RKKLISLLGAALKRKLKKRKNQIFGF | Soluble |
| Turingcin-38 | KFLHVLILLTAAALKKAKALAK | Soluble |
| Turingcin-39 | RLLSILAILIFRKSNNVIFINIIINKK | Insoluble |
| Turingcin-40 | KAARRKKLFLAKRIRTIFK | Soluble |
| Turingcin-41 | KRRKMRKKWAKYINFKKITR | Soluble |
| Turingcin-42 | RKKMYIYARSKLRNWVLR | Soluble |
| Turingcin-43 | RLKEKRIIKVRATLRLNNIFTTK | Soluble |
| Turingcin-44 | RRRRSRKKIFVKFIL | Insoluble |
| Turingcin-45 | RKKHVSLIINNRKQLLHNIKFRSKI | Soluble |
| Turingcin-46 | FRWWLYKRALAATTIKIKRRK | Soluble |
| Turingcin-47 | WIRRLIIARWFRTK | Soluble |
| Turingcin-48 | KNIWTHLALKIRLKLNRAE | Soluble |
| Turingcin-49 | RKRVIKPSLALNKWLW | Soluble |
| Turingcin-50 | RVKRLKLYINALTKLV | Soluble |
| Turingcin-51 | TKRRIIKYAAFKFKKALI | Soluble |
| Turingcin-52 | WLLNRILIIHNLAKI | Insoluble |
| Turingcin-53 | KYLTIHFRKTIRFWIRVIK | Soluble |
| Turingcin-54 | RYYLAHVIYIKIWNLQALFKKWKIII | Insoluble |
| Turingcin-55 | RSWIISLIRRL | Soluble |
| Turingcin-56 | RLRRRYKRILIQDKRLNKILYLI | Soluble |
| Turingcin-57 | WRHKSLWIRKYLKNLALLA | Soluble |
| Turingcin-58 | KRRNYIIIRLAKNIINKIINNLQ | Soluble |
| Turingcin-59 | KLRRLNKINLINKTILEKQRKINKL | Soluble |
| Turingcin-60 | KKLRIRTMLKRWKWELAFH | Soluble |

**Table S2.** Cytotoxicity [$CC_{50}$ for human keratinocytes (HaCaT), PMA-differentiated human monocytes (THP-1), and $HC_{50}$ for human red blood cells (RBCs)] and minimum antimicrobial activity (lowest MIC observed across all tested pathogens) of Turingcins.

| Peptide | $CC_{50}$ (HaCaT) | $CC_{50}$ (THP-1) | $HC_{50}$ (RBCs) | MIC |
|---|---|---|---|---|
| Turingcin-2 | 9.47 | 13.69 | 48.78 | 0.78 |
| Turingcin-3 | 74.82 | 94.92 | 2117 | 3.12 |
| Turingcin-6 | 73.57 | 62.37 | 9527 | 6.25 |
| Turingcin-8 | 7.598 | 23.08 | 5.039 | 1.56 |
| Turingcin-9 | 36.81 | 59.51 | 787.5 | 0.78 |
| Turingcin-11 | 40.87 | 47.74 | 11800 | 3.12 |
| Turingcin-12 | 22.75 | 41.01 | 130.2 | 0.78 |
| Turingcin-13 | 26.7 | 101.4 | 280.1 | 0.78 |
| Turingcin-14 | 29.5 | 116.2 | 207.1 | 12.5 |
| Turingcin-15 | 11.58 | 15.69 | 104.8 | 0.78 |
| Turingcin-16 | 25.2 | 43 | 5520 | 12.5 |
| Turingcin-17 | 15.32 | 37.98 | 2123 | 0.78 |
| Turingcin-18 | 45.39 | 114.1 | 2041 | 6.25 |
| Turingcin-22 | 23.62 | 21.66 | 9.585 | 0.78 |
| Turingcin-23 | 18.73 | 25.43 | 1695 | 1.56 |
| Turingcin-24 | 38.14 | 126.9 | 8461 | 3.12 |
| Turingcin-25 | 22.25 | 21.51 | 328.8 | 0.78 |
| Turingcin-26 | 12.54 | 17.82 | 264.6 | 0.78 |
| Turingcin-27 | 44.39 | 44.65 | 123.4 | 0.78 |
| Turingcin-28 | 32.63 | 116 | 66.13 | 3.12 |
| Turingcin-31 | 39.01 | 170.6 | 3937 | 3.12 |
| Turingcin-32 | 34.84 | 256.8 | 19155 | 50 |
| Turingcin-33 | 25.94 | 152.9 | 4348 | 1.56 |
| Turingcin-34 | 10.68 | 14.27 | 11.63 | 0.78 |
| Turingcin-35 | 84.86 | 196.7 | 357.4 | 1.56 |
| Turingcin-37 | 14.69 | 11.45 | 98.23 | 0.78 |
| Turingcin-38 | 29.68 | 32.43 | 135.7 | 1.56 |
| Turingcin-40 | 63.13 | 104.6 | 4805 | 6.25 |
| Turingcin-41 | 41.91 | 45.15 | 2840 | 6.25 |
| Turingcin-42 | 39.45 | 84.42 | 2403 | 1.56 |
| Turingcin-43 | 12.63 | 314.4 | 1350 | 0.78 |
| Turingcin-45 | 36.7 | 328.9 | 7054 | 12.5 |
| Turingcin-46 | 54.81 | 74.42 | 126.9 | 1.56 |
| Turingcin-47 | 37.35 | 112 | 75.65 | 1.56 |
| Turingcin-49 | 54.2 | 328.6 | 4579 | 12.5 |
| Turingcin-50 | 27.22 | 99.55 | 248.3 | 3.12 |
| Turingcin-51 | 56.05 | 180.1 | 10540 | 6.25 |
| Turingcin-53 | 440 | 4.83E+53 | 58.15 | 3.12 |
| Turingcin-55 | 80.82 | 2158 | 665 | 3.12 |
| Turingcin-56 | 30.24 | 97.93 | 498.1 | 1.56 |
| Turingcin-57 | 22.63 | 43.42 | 86.24 | 0.78 |
| Turingcin-58 | 26.37 | 29.53 | 7.73 | 3.12 |

| | | | | |
|---|---|---|---|---|
| Turingcin-60 | 229.5 | 105.5 | 1169 | 0.78 |

**Table S3.** Sequence of Turingcin-46 and 18 single-substitution derivatives created by replacing each amino acid with alanine.

| **Peptide** | **Sequence** | **Solubility in autoclaved deionized water** |
|---|---|---|
| Turingcin-46 | FRWWLYKRALAATTIKIKRRK | Soluble |
| F1A-Turingcin-46 | ARWWLYKRALAATTIKIKRRK | Soluble |
| R2A-Turingcin-46 | FAWWLYKRALAATTIKIKRRK | Soluble |
| W3A-Turingcin-46 | FRAWLYKRALAATTIKIKRRK | Soluble |
| W4A-Turingcin-46 | FRWALYKRALAATTIKIKRRK | Soluble |
| L5A-Turingcin-46 | FRWWAYKRALAATTIKIKRRK | Soluble |
| Y6A-Turingcin-46 | FRWWLAKRALAATTIKIKRRK | Soluble |
| K7A-Turingcin-46 | FRWWLYARALAATTIKIKRRK | Soluble |
| R8A-Turingcin-46 | FRWWLYKAALAATTIKIKRRK | Soluble |
| L10A-Turingcin-46 | FRWWLYKRAAAATTIKIKRRK | Soluble |
| T13A-Turingcin-46 | FRWWLYKRALAAATIKIKRRK | Soluble |
| T14A-Turingcin-46 | FRWWLYKRALAATAIKIKRRK | Soluble |
| I15A-Turingcin-46 | FRWWLYKRALAATTAKIKRRK | Soluble |
| K16A-Turingcin-46 | FRWWLYKRALAATTIAIKRRK | Soluble |
| I17A-Turingcin-46 | FRWWLYKRALAATTIKAKRRK | Soluble |
| K18A-Turingcin-46 | FRWWLYKRALAATTIKIARRK | Soluble |
| R19A-Turingcin-46 | FRWWLYKRALAATTIKIKARK | Soluble |
| R20A-Turingcin-46 | FRWWLYKRALAATTIKIKRAK | Soluble |
| K21A-Turingcin-46 | FRWWLYKRALAATTIKIKRRA | Soluble |

**Table S4.** Physicochemical properties of Turingcin-46 and 18 single-substitution derivatives obtained from the DBAASP server [2]. Note that Eisenberg and Weiss scale [3] was chosen as the hydrophobicity scale.

| Peptide | Normalized Hydrophobic Moment | Normalized Hydrophobicity | Net Charge | Isoelectric Point | Penetration Depth | Tilt Angle | Disordered Conformation Propensity | Linear Moment | Propensity to in vitro Aggregation | Angle Subtended by the Hydrophobic Residues | Amphiphilicity Index | Propensity to PPII coil |
|---|---|---|---|---|---|---|---|---|---|---|---|---|
| Turingcin-46 | 0.18 | -0.17 | 8 | 12.15 | 24 | 22 | -0.34 | 0.28 | 1.57 | 60 | 2.07 | 0.98 |
| F1A-Turingcin-46 | 0.13 | -0.07 | 8 | 12.16 | 23 | 28 | -0.35 | 0.28 | 1.57 | 60 | 2.07 | 0.97 |
| R2A-Turingcin-46 | 0.31 | -0.35 | 7 | 11.94 | 21 | 23 | -0.19 | 0.36 | 82.9 | 70 | 1.95 | 0.97 |
| W3A-Turingcin-46 | 0.2 | -0.16 | 8 | 12.15 | 24 | 25 | -0.3 | 0.28 | 1.57 | 60 | 1.74 | 0.97 |
| W4A-Turingcin-46 | 0.18 | -0.16 | 8 | 12.15 | 24 | 23 | -0.3 | 0.28 | 1.57 | 60 | 1.74 | 0.97 |
| L5A-Turingcin-46 | 0.1 | -0.09 | 8 | 12.15 | 25 | 22 | -0.36 | 0.27 | 1.56 | 60 | 2.07 | 0.98 |
| Y6A-Turingcin-46 | 0.19 | -0.12 | 8 | 12.6 | 24 | 26 | -0.29 | 0.28 | 1.56 | 60 | 1.83 | 0.96 |
| K7A-Turingcin-46 | 0.1 | -0.43 | 7 | 12.15 | 19 | 53 | -0.2 | 0.35 | 36.81 | 60 | 1.89 | 0.98 |
| R8A-Turingcin-46 | 0.3 | -0.35 | 7 | 11.94 | 22 | 37 | -0.19 | 0.31 | 6.19 | 90 | 1.95 | 0.97 |
| L10A-Turingcin-46 | 0.25 | -0.09 | 8 | 12.15 | 24 | 23 | -0.36 | 0.29 | 0 | 60 | 2.07 | 0.98 |
| T13A-Turingcin-46 | 0.22 | -0.26 | 8 | 12.15 | 24 | 22 | -0.3 | 0.28 | 3.91 | 80 | 2.07 | 0.97 |
| T14A-Turingcin-46 | 0.09 | -0.26 | 8 | 12.15 | 24 | 22 | -0.3 | 0.28 | 3.93 | 60 | 2.07 | 0.97 |
| I15A-Turingcin-46 | 0.19 | -0.1 | 8 | 12.15 | 24 | 22 | -0.38 | 0.3 | 0 | 60 | 2.07 | 0.97 |
| K16A-Turingcin-46 | 0.43 | -0.43 | 7 | 12.15 | 23 | 32 | -0.2 | 0.33 | 29.89 | 80 | 1.89 | 0.98 |
| I17A-Turingcin-46 | 0.22 | -0.1 | 8 | 12.15 | 24 | 22 | -0.38 | 0.31 | 1.56 | 60 | 2.07 | 0.97 |
| K18A-Turingcin-46 | 0.15 | -0.43 | 7 | 12.15 | 14 | 85 | -0.2 | 0.23 | 1.57 | 80 | 1.89 | 0.98 |
| R19A-Turingcin-46 | 0.33 | -0.35 | 7 | 11.94 | 23 | 31 | -0.19 | 0.24 | 1.57 | 60 | 1.95 | 0.97 |
| R20A-Turingcin-46 | 0.31 | -0.35 | 7 | 11.94 | 24 | 22 | -0.19 | 0.23 | 1.57 | 60 | 1.95 | 0.97 |
| K21A-Turingcin-46 | 0.12 | -0.43 | 7 | 12.15 | 23 | 31 | -0.2 | 0.25 | 1.57 | 80 | 1.89 | 0.98 |
